# A conserved glutamate residue in RPM1-interacting protein4 is ADP-ribosylated by *Pseudomonas* effector AvrRpm2 to activate RPM1-mediated response

**DOI:** 10.1101/2021.09.01.458589

**Authors:** Minsoo Yoon, Martin Middleditch, Erik Rikkerink

## Abstract

Gram-negative bacterial plant pathogens directly inject effectors into their hosts to hijack and manipulate metabolism, eluding the frontier surveillance at the cell surface. The effector AvrRpm1_Pma_ from *Pseudomonas syringae* pv. *maculicola* functions as an ADP-ribosyl transferase, modifying RPM1-interacting protein4 (RIN4), leading to the activation of Arabidopsis resistance protein RPM1. We identified the ADP-ribosyl transferase activity of another bacterial effector AvrRpm2_Psa_ from *Pseudomonas syringae* pv. *actinidiae* via infection using a *Pseudomonas syringae* pv. *tomato* strain following Agrobacterium-mediated transient expression of RIN4 in *N. benthamiana*. We conducted mutational analysis in combination with mass spectrometry to genetically locate the modified residue. We show that a conserved glutamate residue (E156) of AtRIN4 is the target site for AvrRpm2_Psa_ by demonstrating the modified AtRIN4 with E156A substitution is no longer ADP-ribosylated. Accordingly, naturally occurring soybean and snap bean RIN4 homologs with no glutamate at the positions corresponding to the E156 of AtRIN4 are not ADP-ribosylated by AvrRpm2_Psa_. In contrast with another effector AvrB, modifications of potential phosphorylation sites including T166 in AtRIN4 affected neither ADP-ribosylation nor RPM1 activation by AvrRpm2_Psa_. This study suggests that separate biochemical reactions by different pathogen effectors may trigger the activation of the same resistance protein through distinct modifications of RIN4.

**One sentence summary:** A conserved glutamate residue (E156) in the C-NOI domain of RPM1-interacting protein4 is ADP-ribosylated by *Pseudomonas* effector AvrRpm2 to activate RPM1-mediated defence response, independently of phosphorylation at T166.

## INTRODUCTION

Bacterial plant pathogens such as *Pseudomonas, Ralstonia, Xanthomonas*, and *Erwinia* can cause a variety of diseases in economically important crop plants (Mansfield et al., 2012). Over long periods of co-existence, both hosts and pathogens have evolved features to combat each other (Bent and Mackey, 2007). At the battle frontier on the cell surface (Malinovsky et al., 2014), various pattern recognition receptors (Macho and Zipfel, 2015; Kong et al., 2021) sense the presence of a pathogen by recognising pathogen-associated molecular patterns (PAMPs) (Ingle et al., 2006), which are signature molecules of pathogens, for example, flagellin or lipopolysaccharides (Felix et al., 1999; Yu et al., 2021), also known as microbe-associated molecular patterns (Ausubel, 2005). Evading host surveillance at the cell wall, pathogens directly inject bacterial proteins into plant cells via a specialised delivery apparatus known as the type III secretion system (T3SS) (Coombes, 2009; Buttner, 2012; Puhar and Sansonetti, 2014). Successful proliferation of pathogens depends on secreted proteins known as type III effectors (T3Es) (Alfano and Collmer, 2004; Block et al., 2008; Block and Alfano, 2011).

Unlike PAMPs, which are fragments from bacterial molecules often conserved for essential microbial life, effectors are specially designed tools for aggressive host colonisation. Effectors may have specific enzymatic activities such as phosphorylation, ubiquitination, acetylation, proteolysis, or ADP-ribosylation (Ribet and Cossart, 2010). Once translocated inside plant cells, T3Es manipulate essential cellular processes to promote pathogen virulence and neutralize defence responses. However, this effector injection strategy is counterbalanced by another layer of host responses orchestrated by resistance (R) genes (Belkhadir et al., 2004; DeYoung and Innes, 2006). Once activated, an R gene may lead to strong defence responses including a hypersensitive responses (HR), often manifested by localised cell death to minimise the spread of infection (Guo et al., 2009; Lindeberg et al., 2012). The encoded R proteins may recognise translocated pathogen effectors via direct physical association or indirectly through perception of the enzymatic activities required for effector function (Dangl and Jones, 2001; van der Hoorn and Kamoun, 2008).

The T3E AvrRpm1_Pma_ from the phytopathogen *Pseudomonas syringae* pv. *maculicola* (Pma) triggers activation of Arabidopsis resistance protein RPM1 (Mackey et al., 2002). The RPM1-mediated defence response is closely linked to the modification of RPM1-interacting protein4 (RIN4) (Chung et al., 2011; Liu et al., 2011; Xu et al., 2017). All events involved in the sequence of interactions, translocation of the bacterial effector into the host via T3SS, posttranslational modification of the host protein RIN4 (Kim et al., 2005), and the activation of resistance protein RPM1 for eventual defence response, are critical for the progression of resistance or disease (Dodds and Rathjen, 2010; Cui et al., 2015). In particular, the ubiquitous plant protein RIN4, a probable regulator of plant immunity (Rikkerink, 2018; Toruno et al., 2019), is a target for multiple effectors (Mackey et al., 2003; Chung et al., 2014; Zhao et al., 2021). However, the physiological or functional role of RIN4 is not clearly defined despite the wide distribution of this protein among plants including mosses (Afzal et al., 2013). The bacterial effector AvrRpm1_Pma_ functions as an ADP-ribosyl transferase, modifying RIN4 (Cherkis et al., 2012; Redditt et al., 2019). The target residues of AvrRpm1_Pma_ were identified via mass spectrometry analysis following Agrobacterium-mediated transient expression of a soybean RIN4 homolog GmRIN4b in *N. benthamiana*. It was reported that a D153 substitution in the C-terminal nitrate-induced (NOI) domain of AtRIN4 inhibited phosphorylation of T166 and eventually inhibited the RPM1-mediated restriction of pathogen growth, supposedly by blocking ADP-ribosylation at this position (Redditt et al., 2019). However, it is still not clear how the ADP-ribosyl transferase activity of AvrRpm1_Pma_, the phosphorylation of the RIN4, and the activation of RPM1 are interconnected to initiate the host response.

Kiwifruit canker disease (Vanneste et al., 2013; Donati et al., 2020) caused by *Pseudomonas syringae* pv. *actinidiae* (Psa) was first reported in Japan (Serizawa et al., 1989) followed by China, Korea, Italy, and then it rapidly spread around the world (Fang et al., 1990; Koh et al., 1994; Scortichini, 1994; McCann et al., 2017). Psa strains collectively have about 50 T3E loci (McCann et al., 2013). While most Psa effectors have not been functionally characterised, some effectors show varying degrees of sequence similarities with previously characterised effectors from other related species such as *Pseudomonas syringae* pv. *tomato* (HopQ1_Pto_ or HopF2_Pto_), *Pseudomonas syringae* pv. *syringae* (AvrRpm1_Psy_ or HopZ3_Psy_), or *Pseudomonas syringae* pv. *maculicola* (AvrRpm1_Pma_), raising the possibility that some effector functions are conserved at least partially, even though the pathogens have different hosts (Cunnac et al., 2009; Baltrus et al., 2011; Dharmaraj, 2018). In particular, most Psa strains have AvrRpm2_Psa_ loci, having about 50% protein sequence identity with AvrRpm1_Pma_, in various allelic forms (McCann et al., 2013; Fujikawa and Sawada, 2016), suggesting frequent selection and counter-selection during host-pathogen interactions.

ADP-ribosylation is a reversible posttranslational modification (PTM) catalysed by a group of enzymes known as ADP-ribosyl transferases (ARTs) (Hottiger et al., 2010). They are sub-classified based on conserved motifs of either H-Y-E (Diphteria toxin or DTX family) or R-S-E (Cholera toxin or CTX family) in the catalytic domains (Simon et al., 2014; Mikolcevic et al., 2021). Recent studies show that ADP-ribosylation is exploited both by bacteria to achieve stealth attacks on their hosts and by plants to launch effective defence (Feng et al., 2016). Mass-spectrometry analysis has become the main analytical tool for identifying PTMs (Doll and Burlingame, 2015). However, there is a major technical limitation of mass spectrometry analysis in locating target sites of ADP-ribosylation. Due to the lability of the bond between the ADP-ribose moiety and the side-chain of the modified amino acid during fragmentation, compared with the relatively stable peptide bonds between amino acid residues, generating all possible combinations of fragment ions retaining the ADP-ribose moiety is often difficult (Rosenthal et al., 2015; Hendriks et al., 2019). Combined with the fact that many different amino acid residues can be modified (Cohen and Chang, 2018), accurate localization of ADP-ribosylation can be analytically challenging.

Here we report the identification of the target residue in RIN4 for the bacterial effector AvrRpm2_Psa_, which also functions as an ADP-ribosyl transferase. To complement the limitation of mass spectrometry, which eventually could not generate all necessary combinations of fragment ions required for unambiguous identification of the modified residue, we conducted mutational analysis to genetically identify the target. To preclude potentially aberrant activities of the bacterial effectors expressed *in planta*, we delivered the bacterial protein via infection using a Pto strain instead of Agrobacterium-mediated transient expression. With the combination of mass spectrometry, mutational analysis, and infection delivery, we located a conserved glutamate residue (E156) in AtRIN4 as the target for the AvrRpm2_Psa_ activity. We found matching polymorphisms at positions corresponding to E156 of AtRIN4 with the RPM1-mediated HR phenotypes among naturally occurring RIN4 homologs of soybean (*Glycine max*), snap bean (*Phaseolus vulgaris*) and apple (*Malus* x *domestica*), demonstrating that the RIN4 target site for this effector is conserved across plant species.

## RESULTS AND DISCUSSION

### AvrRpm2_Psa_ functions as an ADP-ribosyl transferase modifying RIN4

Sequence alignment with known ADP-ribosyl transferases such as diphtheria toxin or Exotoxin A (Exo T) suggested that AvrRpm1_Pma_ has three conserved residues (H63, Y122, and D185) in catalytic domains (Cherkis et al., 2012). It was later biochemically shown that AvrRpm1_Pma_ functions as an ADP-ribosyl transferase to modify RIN4, leading to the activation of Arabidopsis resistance protein RPM1 (Redditt et al., 2019). AvrRpm1_Pma_ has the H-Y-D motif in potential catalytic domains and therefore may be classified as a member of the DTX family along with *Pseudomonas aeruginosa* Exo T, a T3E required for full virulence in animal model of an acute pneumonia, which has the H-Y-E motif (Garrity-Ryan et al., 2004). While AvrRpm2_Psa_ shares about 50% protein sequence with AvrRpm1_Pma_, the proposed three critical residues (H-Y-D) in AvrRpm1_Pma_ are also conserved in AvrRpm2_Psa_ (Figure 1A).

**Figure 1.**
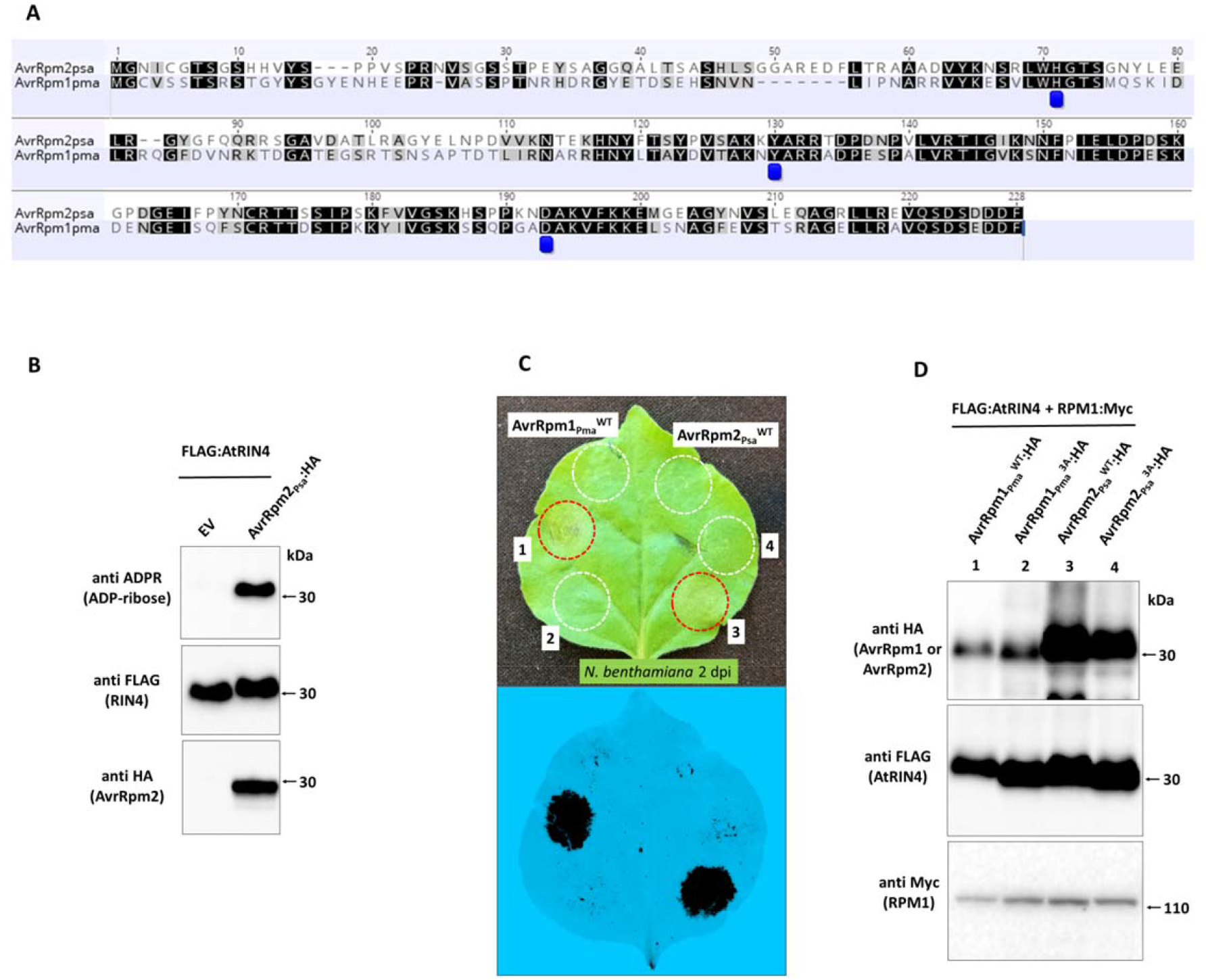
Transient co-expression of AtRIN4 and AvrRpm2_Psa_ (biovar 5) in *N. benthamiana* via Agrobacterium. **A**. Sequence alignment of AvrRpm2_Psa_ and AvrRpm1_Pma_. The residues (H63, Y122, and D185) in the conserved H-Y-D motif of AvrRpm1_Pma,_ proposed previously (Cherkis et al., 2012), are marked with blue labels. The corresponding residues in the H-Y-D motif of AvrRpm2_Psa_ are H68, Y125, and D188. **B**. Western blot analysis demonstrating the ADP-ribosyl transferase activity of AvrRpm2_Psa_. Proteins were extracted from *N. benthamiana* leaves co-expressing AvrRpm2_Psa_:HA and FLAG:AtRIN4 via Agrobacterium at 2 d post infiltration. ADP-ribosylated proteins were detected using the anti ADPR binding reagent. FLAG:RIN4 and AvrRpm2_Psa_:HA were detected using corresponding antibodies. **C**. RPM1-mediated HR (hypersensitive response) assay. An *N. benthamiana* leaf co-expressing FLAG:AtRIN4 and RPM1:Myc with either AvrRpm1_Pma_ or AvrRpm1_Pma_ in different combinations (1: AvrRpm1_Pma_^WT^:HA (wild type); 2: AvrRpm1_Pma_^3A^ (triple substitutions: H63A, Y122A, D185A); 3: AvrRpm2_Psa_^WT^:HA (wild type); 4: AvrRpm2_Psa_^3A^ (triple substitutions: H68A, Y125A, D188A) delivered via Agrobacterium. Red circles denote HR and white circles denote no HR. The fluorescence of the same leaf monitored under a 488 nm tray in the ChemiDoc™ is also shown (bottom). Images were taken 2 d post Agrobacterium infiltration. **D**. Western blot analysis using proteins extracted from the corresponding areas of the leaf used in **C** 2 d post Agrobacterium injection.

We tested AvrRpm2_Psa_ for its function as an ADP-ribosyl transferase. Proteins were extracted from *N. benthamiana* leaves transiently co-expressing AtRIN4 and AvrRpm2_Psa_ via Agrobacterium. When Western blot analysis was performed using anti-ADPR binding reagent to detect ADP-ribosylated proteins, the AtRIN4 protein band was detected, demonstrating that AvrRpm2_Psa_ functions as an ADP-ribosyl transferase modifying AtRIN4 (Figure 1B). When AvrRpm2_Psa_ was co-expressed with AtRIN4 and RPM1 via Agrobacterium in *N. benthamiana*, an RPM1-mediated HR was observed (Figure 1C, top). When the fluorescence from the infiltrated leaf was monitored in the ChemiDoc™ with trays specific for 488nm, loss of green specks due to cell collapse were more clearly visible (Figure 1C, bottom) (Yoon and Rikkerink, 2020). As reported previously, modifications of the three sites (H-Y-D) in AvrRpm1_Pma_ (AvrRpm1_Pma_^3A^ with triple substitutions at H63A, Y122A, and D185A) resulted in the loss of the RPM1-mediated HR (Cherkis et al., 2012; Redditt et al., 2019). Modifications of the three corresponding residues in AvrRpm2_Psa_ (AvrRpm2_Psa_^3A^ with the corresponding triple substitutions at H68A, Y125A, and D188A) similarly resulted in the loss of the RPM1-mediated response, demonstrating the importance of the H-Y-D motif in the two bacterial effectors. Western blot analysis (Figure 1D) showed that the modified proteins accumulated similarly when compared with their corresponding wild type proteins, suggesting the observed differences in the HR are due to the changes in their biochemical activities, not in their accumulation.

RIN4 is a widely-distributed plant protein capable of physically associating with other proteins, making multi-protein complexes (Sun et al., 2014; Rikkerink, 2018; Ray et al., 2019). There are several bacterial effectors known to physically associate with RIN4, including AvrB (Lee et al., 2004), AvrRpm1_Pma_ (Mackey et al., 2002), AvrRpm1_Psa_ (Yoon and Rikkerink, 2020), HopF2_Pto_ (Wilton et al., 2010), and HopZ3_Psy_ (Lee et al., 2015b), suggesting frequent participation of RIN4 in pathogen-plant interactions. To test the protein association with RIN4, AvrRpm2_Psa_ with an epitope tag (AvrRpm2_Psa_:HA) was co-expressed with AtRIN4, tagged with a different epitope (FLAG:AtRIN4), in *N. benthamiana* via Agrobacterium. Both proteins were detected in the Western blot analysis with corresponding antibodies (Figure 2A). Proteins were immunoprecipitated with anti HA-conjugated magnetic beads to isolate AvrRpm2_Psa_. Western blot analysis showed that AtRIN4 was co-precipitated with AvrRpm2_Psa_, showing AtRIN4 physically associated with AvrRpm2_Psa_ (Figure 2A, third lane, Co-IP panel). There was no comparable AtRIN4 co-precipitated with the empty vector (EV) or HA-tagged AtRIN4 (HA:AtRIN4), showing the protein-protein interaction between AvrRpm2_Psa_ and AtRIN4 was specific. AtRIN4 co-expressed with AvrRpm2_Psa_ showed a change in mobility during electrophoresis. As shown in the Western blot analysis, AtRIN4 co-expressed with AvrRpm2_Psa_ (Figure 2B, even-numbered) showed a consistently slower migration in SDS-PAGE compared with those from leaves without AvrRpm2_Psa_ (odd-numbered), suggesting a change in the structure of the ADP-ribosylated RIN4 compared with the unmodified RIN4. Similar changes in the mobility of RIN4 during electrophoresis were reported previously with co-expressed AvrRpm1_Pma_ (Redditt et al., 2019).

**Figure 2.**
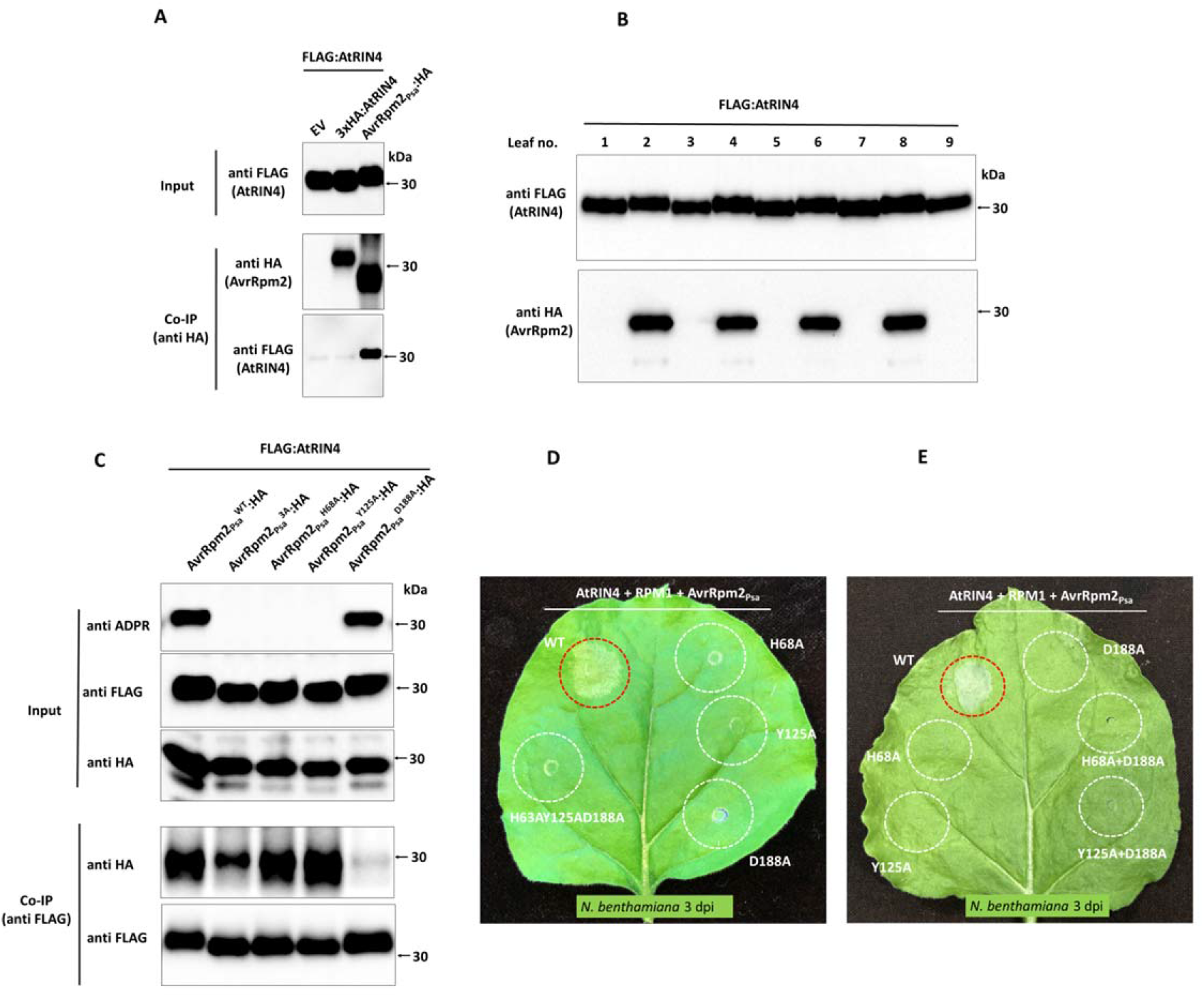
Western blot analysis of AtRIN4 transiently co-expressed with AvrRpm2_Psa_ in *N. benthamiana* via Agrobacterium. **A**. Proteins were extracted from *N. benthamiana* leaves co-expressing FLAG:AtRIN4 with either Empty vector (lane 1), HA:AtRIN4 (lane 2), or AvrRpm2_Psa_:HA (lane 3) via Agrobacterium. Protein extracts were immunoprecipitated using anti HA magnetic beads. Precipitated proteins were probed with corresponding antibodies. **B**. Western blot analysis showed the ADP-ribosylated AtRIN4 migrated more slowly compared with unmodified AtRIN4 during SDS-PAGE. Protein extracts from *N. benthamiana* leaves co-expressing FLAG:AtRIN4 with AvrRpm2_Psa_:HA (even-numbered) or expressing only FLAG:AtRIN4 (odd-numbered) were probed. **C**. Western blot analysis of different AvrRpm2_Psa_ alelles (AvrRpm2_Psa_^WT^, AvrRpm2_Psa_^H68A^, AvrRpm2_Psa_^Y125A^, AvrRpm2_Psa_^H188A^, and AvrRpm2_Psa_^3A^) co-expressed with AtRIN4 via Agrobacterium (Input panel). For Co-IP of modified AvrRpm2 with AtRIN4 (Co-IP panels), protein extracts were precipitated using anti FLAG magnetic beads to isolate FLAG:AtRIN4. Precipitated proteins were probed using corresponding antibodies. **D**. Modified AvrRpm2 alleles (AvrRpm2_Psa_^H68A^, AvrRpm2 _Psa_^Y125A^, and AvrRpm2_Psa_^D188A^) were co-expressed with RPM1 and AtRIN4 on an *N. benthamiana* leaf to assess the corresponding RPM1-mediated HR. **E**. Combinations of modified AvrRpm2 alleles (AvrRpm2_Psa_^D188A^ with AvrRpm2_Psa_^H68A^ and AvrRpm2_Psa_^D188A^ with AvrRpm2_Psa_^Y125A^) were co-expressed with RPM1 and AtRIN4 on an *N. benthamiana* leaf. Red circles denote HR and white circles denote no HR. Images were taken at 3 d post Agrobacterium injection (**D** and **E**)

### Modifications in the H-Y-D motif of AvrRpm2_Psa_ affect activity or affinity

Modified AvrRpm1_Pma_ with either H63A, Y122A, or D185A substitution resulted in significant loss in the RPM1-mediated restriction of pathogen growth in Arabidopsis (Cherkis et al., 2012). We made corresponding substitutions of H68A, Y125A, and D188A in the H-Y-D motif of AvrRpm2_Psa_ to create AvrRpm2_Psa_^H68A^, AvrRpm2_Psa_^Y128A^, and AvrRpm2_Psa_^D188A^ to analyse their activities as well as their abilities to physically associate with AtRIN4. Proteins were extracted from *N. benthmiana* leaves co-expressing the modified AvrRpm2_Psa_ with AtRIN4 via Agrobacterium. Western blot analysis showed that AvrRpm2_Psa_^H68A^, AvrRpm2_Psa_^Y125A^ lost the catalytic activity to modify AtRIN4 while AvrRpm2_Psa_^D188A^ retained the activity (Figure 2C, anti ADPR in Input panel). In contrast, Co-IP analysis showed that AvrRpm2_Psa_^D188A^ lost the physical association with AtRIN4 while AvrRpm2_Psa_^H68A^ and AvrRpm2_Psa_^Y125A^ still associated with RIN4 (Figure 2C, anti HA in Co-IP panel). When the modified AvrRpm2_Psa_ proteins were co-expressed with AtRIN4 and RPM1 in *N. benthamiana* via Agrobacterium, all three individual substitutions (H68A, Y125A, or D188A) resulted in the loss of the corresponding RPM1-mediated HR (Fig 2D). Therefore, for the bacterial effector to activate RPM1, the ability to physically associate with RIN4 is also required in addition to the functional ability to modify RIN4. Next, we co-expressed AvrRpm2_Psa_^H68A^ and AvrRpm2_Psa_^D188A^ in *N. benthamiana* with AtRIN4 and RPM1 to see whether RPM1 can be activated by the catalytic activity of AvrRpm2_Psa_^D188A^ to modify AtRIN4 while the protein association with AtRIN4 can be separately provided by AvrRpm2_Psa_^H68A^. We also similarly tested the combined expression of AvrRpm2_Psa_^Y125A^ with AvrRpm2_Psa_^D188A^. Neither combination triggered the corresponding RPM1-mediated HR in *N. benthamiana*, demonstrating that the protein association with AtRIN4 and the catalytic activity to modify AtRIN4 cannot be separately provided to activate RPM1 (Figure 2E).

### Mass spectrometry analysis of AtRIN4 co-expressed with AvrRpm1_Pma_ in *N. benthamiana* via Agrobacterium

Previously, two residues N12 and D185 in GmRIN4b were identified as the target sites for AvrRpm1_Pma_ (Redditt et al., 2019). To locate the target residue(s) in AtRIN4 for the AvrRpm2_Psa_ activity, we adopted a similar approach. As a preliminary control analysis we first analysed the ADP-ribosylation of AtRIN4 with co-expressed AvrRpm1_Pma_. An N-FLAG tagged AtRIN4 (FLAG:AtRIN4) was co-expressed with AvrRpm1_Pma_ in *N. benthamiana* via Agrobacterium and isolated by immunoprecipitation using anti FLAG-conjugated magnetic beads (Figure S1A). The protein band corresponding to AtRIN4 was excised from the gel and confirmed by Western blot analysis (Figure S1B). After LC-MS/MS analysis of the excised AtRIN4 protein band, we detected ADP-ribosylated peptides from the two nitrate-induced (NOI) domains. Available fragment evidence suggested that two residues (N157 and N158) in the C-NOI domain and the corresponding two residues (E16 and N17) in the N-NOI domian were likely to be ADP-ribosylated. Not every possible combination of fragment ions were recovered, and most ADP-ribose moieties identified were in degraded forms such as phospho-ribose or ribose, suggesting the lability of the ADP-ribose moiety during fragmentation. Examples of fragmentation data are shown in Figure S1C and Figure S1D.

The target sites of AvrRpm1_Pma_ previously identified in GmRIN4b (N12 and D185) correspond to N11 and D153 in AtRIN4, respectively (Redditt et al., 2019), which are clearly distinct from the target residues we identified (E16, N17, N157, N158) by directly analysing AtRIN4. In an effort to resolve the discrepancy, we created modified RIN4 alleles by replacing target sites identified by both analyses with alanine to prevent modification on these sites. GmRIN4^N12AD185A^ and corresponding AtRIN4^N11AD153A^ were created based on the earlier analysis of GmRIN4b, and AtRIN4^E16AN17AN157AN158A^ was created based on our mass spectrometry analysis. Proteins were extracted from *N. benthamiana* leaves co-expressing AvrRpm1_Pma_ with the modified RIN4 proteins via Agrobacterium. Western blot analysis showed that all of the modified RIN4 proteins were still ADP-ribosylated (Figure S2A and Figure S2B) and also activated RPM1 (Figure S2C). Even when further modification with all six residues (N11, E16, N17, D153, N157, and N158) collectively identified by both analyses were replaced, the modified protein (AtRIN4^E11AE16AN17AD153AN157AN158A^) was still ADP-ribosylated when co-expressed with AvrRpm1_Pma_ (Figure S2B).

The absence of the target residues identified by the mass spectrometry analyses affected neither the ADP-ribosylation by AvrRpm1_Pma_ nor the activation of RPM1. Therefore, the identification of target residues may have been incorrect with the wrong residues identified due to the hyperactivity of the bacterial effector expressed *in planta* via Agrobacterium, or the mass spectrometry analyses were incomplete with other target residues still unidentified due to the lability of the bonds between ADP-ribose moieties and amino acids side chains. Another possibility is that there are preferences for AvrRpm1_Pma_ among multiple residues and that some sites are modified only when more preferred sites are absent. We similarly analysed AtRIN4 co-expressed with AvrRpm2_Psa_ in *N. benthamiana* via Agrobacterium. Compared with AvrRpm1_Pma_, fewer ADP-ribosylated peptides were identified with AvrRpm2_Psa_ (Figure S3A) and the intensities of fragment ions were lower in the MS/MS spectra (Figure S3B). Available fragmentation spectra suggested that the target site was likely to be in the C-NOI domain. However, not all combinations of fragment ions were generated and unambiguous identification of the target site was difficult.

### Bacterial effectors can be efficiently delivered into *N. benthamiana* by Pto DC3000Q^-^

Even though a few modified residues were identified, we still could not biochemically verify them as real targets (Figure S2). In particular, it cannot be ruled out that the bacterial effectors expressed *in planta* via Agrobacterium may not have identical properties to the secreted effector via T3SS during an infection. Therefore, we compared the activities of T3SS-delivered effectors with those of the effectors expressed *in planta* via Agrobacterium. Previously, we developed a pathogen assay system in *N. benthamiana* by co-injecting Agrobacterium and the Pto DC3000 strain without hopQ1 (Wei et al., 2007), Pto DC3000Q^-^, simultaneously (Yoon and Rikkerink 2020). At low bacterial concentrations (OD_600_=0.00001 to 0.0001 for Pto DC3000Q^-^ and OD_600_=0.01 to 0.04 for Agrobacterium), Agrobacterium and Pto DC3000Q^-^ did not interfere with each other, facilitating bacterial pathogen growth assays. Buscaill et al (2021) reported a similar disease assay based on sequential infection of Pto DC3000Q^-^ following Agrobacterium-mediated transient expression in *N. benthamiana* (Buscaill et al., 2021).

By adopting a similar sequential approach, we first transiently expressed RIN4 in *N. benthamiana* via agrobacterium. At 2 d post infiltration of Agrobacterium, the leaves pre-infiltrated with Agrobacterium were infected with Pto DC3000Q^-^(AvrRpm2_Psa_) at a high bacterial concentration (OD_600_=1.0). There were no significant differences in RIN4 accumulation with Agrobacterium concentrations of OD_600_=0.02 to 0.4. The *N. benthamiana* leaf areas infected with Pto DC3000Q^-^(AvrRpm2_Psa_) showed clear signs of infection at 15 h post infiltration (Figure 3A). At 1 d post infection of Pto DC3000Q^-^(AvrRpm2_Psa_), proteins were extracted from the infected leaves for Western blot analysis. The ADP-ribosylation of RIN4 was clearly detected from a Western blot prepared with the protein extracts without further fractionation or concentration (Figure 3B). In the transgenic Arabidopsis-Pto DC3000 system, expressing high concentration of proteins is difficult because host cells collapse as infections progress. In this Agrobacterium-DC3000Q^-^ system in *N. benthamiana*, Agrobacterium-mediated transient expression prior to pathogen infection ensured high accumulation of RIN4 protein while through infection the expressed bacterial effector was efficiently secreted from the highly concentrated Pto DC3000Q^-^(AvrRpm2_Psa_) into the host, facilitating the detection of the posttranslational modification.

**Figure 3.**
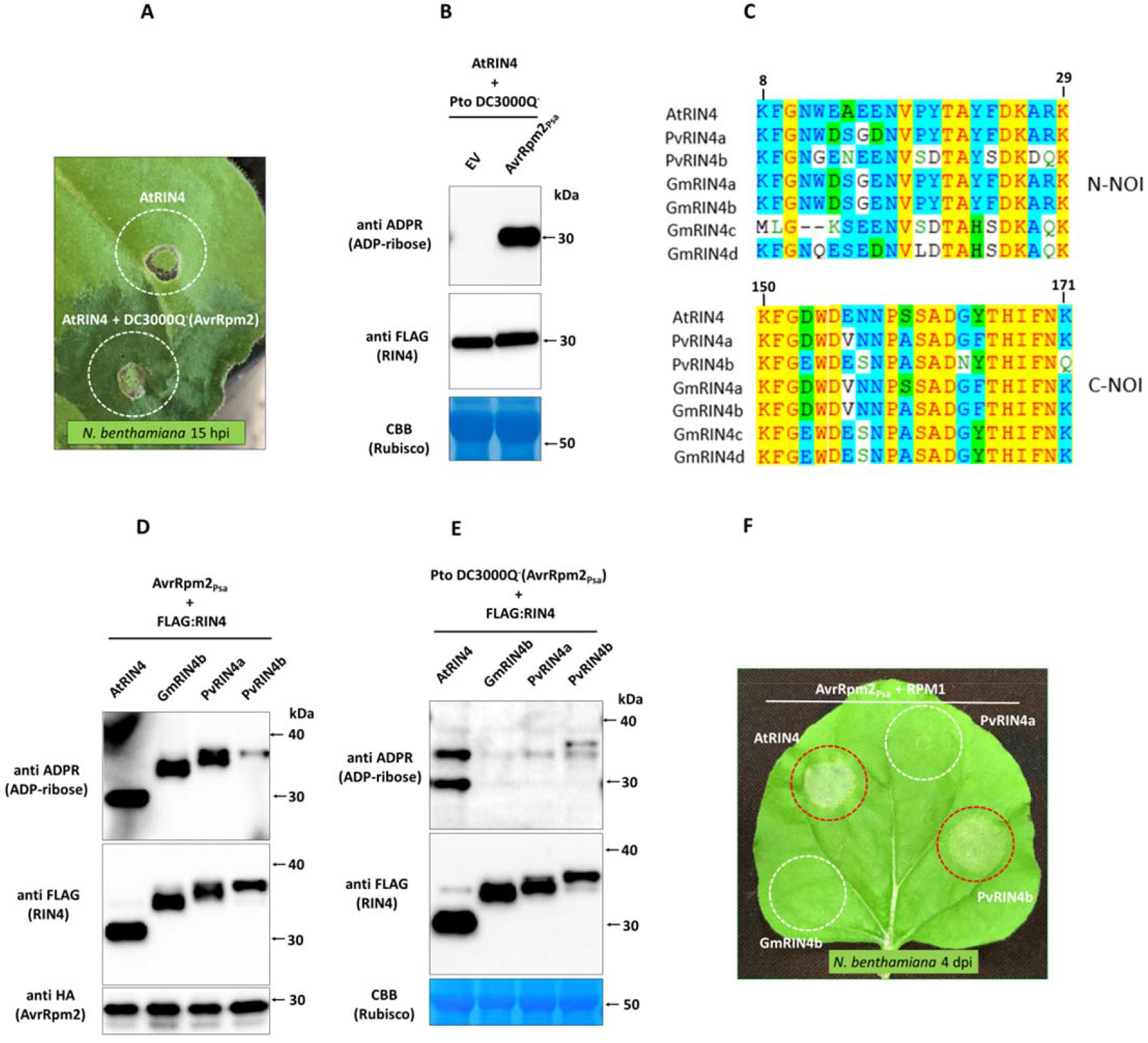
Secretion of bacterial effector via infection of Pto DC3000Q^-^ in *N. benthamiana*. **A**. An *N. benthamiana* leaf was infected with Pto DC3000Q^-^(AvrRpm2_Psa_) at a high concentration (OD_600_=1.0, or 5×10^8^ cfu/mL) in the area pre-infiltrated with Agrobacterium 2 d earlier for transient expression of AtRIN4 (marked with a circle, bottom). Dark discolouration and leaf margin malformations indicate Pto infection. The control area infiltrated only with Agrobacterium is also marked (top). Image was taken at 15 h post infection. **B**. Western blot analysis demonstrated that AtRIN4 was ADP-ribosylated upon infection with Pto DC3000Q^-^(AvrRpm2_Psa_). Proteins were extracted from *N. benthamiana* leaves expressing AtRIN4 via Agrobacterium after infection with Pto DC3000Q^-^ (AvrRpm2_Psa_). ADP-ribosylated proteins were detected using anti ADPR binding reagent. (EV: control Pto DC3000Q^-^ with an empty vector). **C**. Alignment of the two NOI sequences in RIN4 homologs of Arabidopsis (AtRIN4), snap bean (PvRIN4a and PvRIN4b) and soybean (GmRIN4a to GmRIN4d). **D**. Western blot analysis of RIN4 co-expressed with AvrRpm2_Psa_ *in planta* via Agrobacterium. **E**. Western blot analysis of RIN4 homologs infected with Pto DC3000Q^-^(AvrRpm2_Psa_). Proteins were extracted as in (**B**). **F**. The indicated RIN4 homologs were co-expressed with AvrRpm2_Psa_ and RPM1 in different areas of an *N. benthamiana* leaf via Agrobacterium to assess the corresponding RPM1-mediated HR. Red circles denote HR and white circles denote no HR.

### Bacterial effectors expressed *in planta* via Agrobacterium may differ in activity to effectors translocated via T3SS

Among the four soybean RIN4 homologs, GmRIN4a and GmRIN4b were ADP-ribosylated when co-expressed with AvrRpm1_Pma_ while GmRIN4c or GmRIN4d were not modified (Redditt et al., 2019). To compare AvrRpm2_Psa_ expressed *in planta* via Agrobacterium with the bacterial effector secreted during an infection, we analysed the ADP-ribosylation in RIN4 homologs of Arabidopsis, soybean, and snap bean (AtRIN4, GmRIN4b, PvRIN4a, and PvRIN4b). PvRIN4a is more closely related to GmRIN4a and GmRIN4b, while PvRIN4b is more similar to GmRIN4c or GmRIN4d (Figure 3C). Proteins were extracted from *N. benthamiana* leaves co-expressing these RIN4 homologs and AvrRpm2_Psa_ via Agrobacterium. Western blot analysis showed that all four RIN4 proteins were ADP-ribosylated (Figure 3D, anti ADPR). In particular, among the two snap bean RIN4 homologs, PvRIN4a was more strongly ADP-ribosylated compared with PvRIN4b. In the next, proteins were extracted from the leaves infected with Pto DC3000Q^-^ (AvrRpm2_Psa_) following transient expression of RIN4 homologs via Agrobacterium. Western blot analysis showed that only AtRIN4 and PvRIN4b were ADP-ribosylated and GmRIN4b and PvRIN4a were not modified (Figure 3E). Modifications in AtRIN4 or PvRIN4b were not significantly different between the two effector deliveries either by the Agrobacterium-mediated transient expression of AvrRpm2_Psa_ or by the infection of Pto DC3000Q^-^ (AvrRpm2_Psa_).

Interestingly, when the RIN4 homologs were co-expressed with AvrRpm2_Psa_ and RPM1 in *N. benthamiana* via Agrobacterium, only AtRIN4 and PvRIN4b triggered the RPM1-mediated HR (Figure 3F). In contrast, GmRIN4b or PvRIN4a resulted in no comparable HR even though they were also ADP-ribosylated by AvrRpm2_Psa_ (Figure 3D). We reasoned that the ADP-ribosylation in GmRIN4b and PvRIN4a may have been caused by an unusual activity of the bacterial effector expressed *in planta* via Agrobacterium. Such modifications may be artefacts created in our experimental system and may not naturally occur during pathogen infection. We conclude that mass spectrometry analysis of a bacterial effector expressed *in planta* via Agrobacterium may not always lead to a *bona fide* identification of ADP-ribosylation catalysed by the T3SS-delivered bacterial effector during an infection.

### Identification of the AvrRpm2_Psa_ target site by mutational analysis

After carefully studying available mass spectra of AtRIN4 peptides, we hypothesised that the target sites are within the NOI domains, and performed mutational analysis by changing candidate residues in the NOI domain sequences. Based on the results from previous mass spectrometry analyses, we tested AtRIN4^6A^, in which the six residues (N11, E16, N17, D153, N157, and N158) collectively identified by earlier analyses as targets were replaced with alanine (A) to prevent modification on these sites (Figure 4A). Western blot analysis showed that AtRIN4^6A^ was still ADP-ribosylated by Pto DC3000Q^-^(AvrRpm2_Psa_) (Figure 4B). We further replaced four more residues to create AtRIN4^10A^. The modified ten residues were the five amino acids (N11, E13, E15, E16, and N17) in the N-NOI domain and the corresponding five residues (D153, D155, E156, N157, and N158) in the C-NOI domain. Western blot analysis showed that AtRIN4^10A^ was no longer ADP-ribosylated by Pto DC3000Q^-^ (AvrRpm2_Psa_) (Figure 4B). When AtRIN4^10A^ was co-expressed with RPM1 and AvrRpm2_Psa_ in *N. benthamiana* via Agrobacterium, no HR was detected compared with AtRIN4^WT^, suggesting RPM1 was not activated (Figure 4C, white circle). In the next step, we systematically reinstated two original residues at a time in AtRIN4^10A^ (one in the N-NOI domain and the corresponding other in the C-NOI domain) to create five different AtRIN4^8A^ alleles (AtRIN4^8A^_N11D153,_ AtRIN4^8A^_E13D155,_ AtRIN4^8A^_E15E156,_ AtRIN4^8A^_N16N157_, and AtRIN4^8A^_N17N158_). The five AtRIN4^8A^ proteins were transiently expressed in *N. benthamiana* via Agrobacterium and the leaves were infected with Pto DC3000Q^-^(AvrRpm2_Psa_). When proteins were extracted and Western blot analysis was performed, we found that ADP-ribosylation was restored in AtRIN4^8A^_E15E156_, while the other four AtRIN4^8A^ alleles were not modified by AvrRpm2_Psa_ (Figure 4D). Accordingly, when the AtRIN4^8A^ proteins were co-expressed with AvrRpm2_Psa_ and RPM1 in *N. benthamiana* via Agrobacterium, only AtRIN4^8A^_E15E156_ triggered the RPM1-mediated HR, matching the genotype with both the biochemical and physiological phenotypes (Figure 4E).

**Figure 4.**
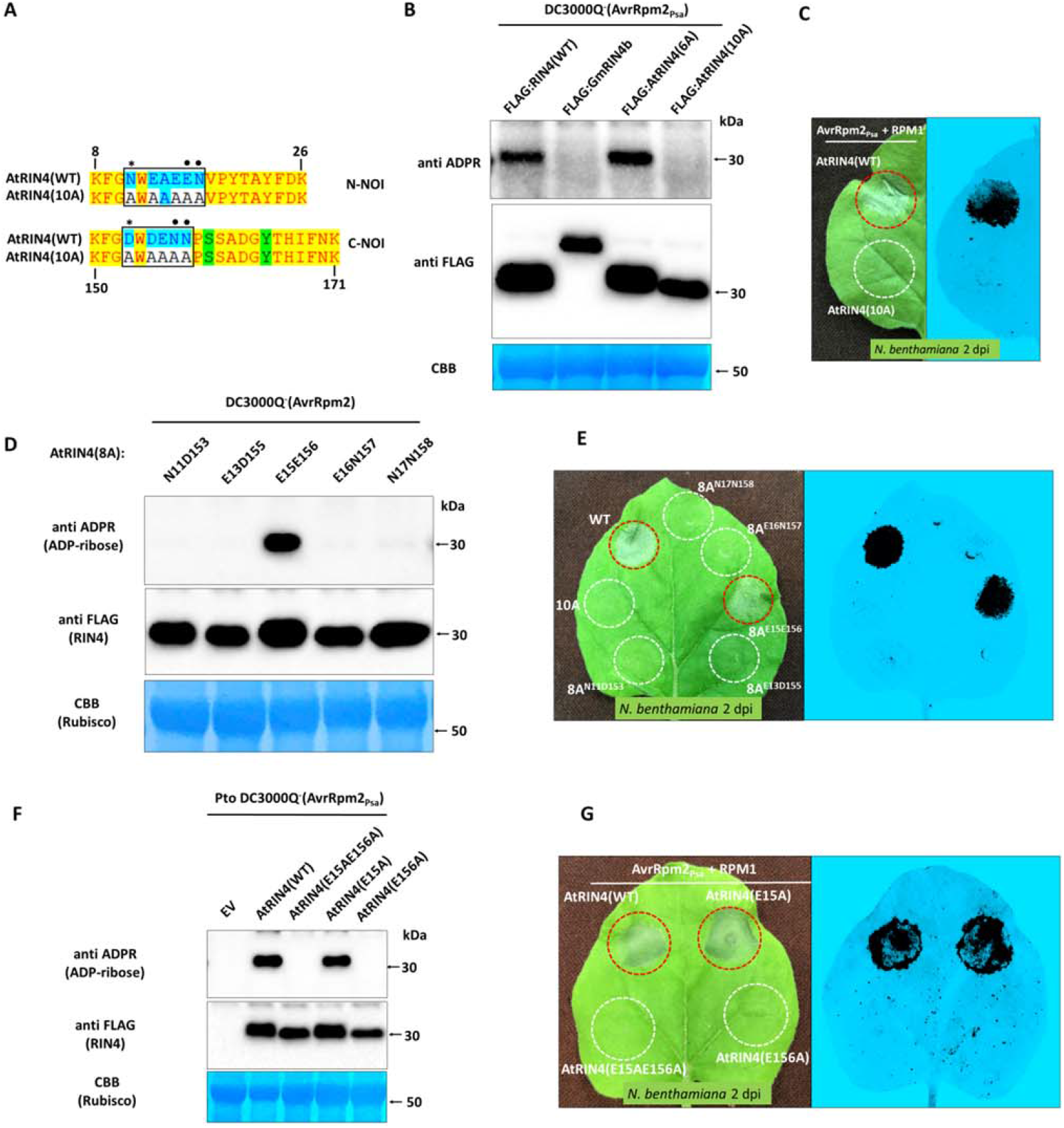
Identification of the target residue for AvrRpm2_Psa_ by mutational analysis. **A**. Modifications generated in the two NOI sequences of AtRIN4 (top). The ten modifications in target amini acids N, D, and E (N11A, E13A, E15A, E16A, N17A, D153A, D155A, E156A, N157A, N158A) in AtRIN4^10A^ are boxed in black (□) and the six modifications in AtRIN4^6A^ (N11A, E16A, N17D, D153A, N157A, N158A) are marked with asterisk (* for the residues N11 and D153 identified as candidates previously) (Redditt et al., 2019) or closed circle (• in E16, N17, N157, and N158 identified as candidates by this study). AtRIN4^10A^ was co-expressed with AvrRpm2_Psa_ and RPM1 in *N. benthamiana*, to assess the RPM1-mediated HR when (bottom). HR was assessed visually or by monitoring fluorescence with ChemiDoc™ at 2 d post Agrobacterium infiltration. **B**. Western blot analysis of RIN4 alleles (AtRIN4^WT^, GmRIN4b^WT^, AtRIN4^6A^ and AtRIN4^10A^) from leaves infected with with Pto DC3000Q^-^(AvrRpm2_Psa_) (left). Proteins were extracted 1 d post infection of Pto DC3000Q^-^(AvrRpm2) in *N. benthamiana* leaves pre-infiltrated with Agrobacterium 2 d earlier for transient expression of RIN4. To assess the RPM1-mediated HR, AtRIN4 proteins were co-expressed with AvrRpm2_Psa_ and RPM1 in *N. benthamiana* (right). The fluorescence image captured in ChemiDoc™ at 2 d post Agrobacterium infiltration is shown. **C**. Western blot analysis of the five AtRIN4^8A^ variant proteins (AtRIN4^8A^_E11D153_, AtRIN4^8A^_E13D155_, AtRIN4^8A^_E15E156_, AtRIN4^8A^_E16N157_, and AtRIN4^8A^_N17N158_) from leaves infected with with Pto DC3000Q^-^ (AvrRpm2_Psa_). Proteins were extracted as in **(B). D**. The five AtRIN4^8A^ proteins were co-expressed with AvrRpm2_Psa_ and RPM1 in *N. benthamiana* via Agrobacterium to assess the corresponding HR. Visual inspection (left) and fluorescence test using the ChemiDoc™ (right) are shown. **E**. Western blot analysis of AtRIN4 proteins (AtRIN4^WT^, AtRIN4^E15AE156A^, AtRIN4^E15A^, and AtRIN4^E156A^) from leaves infected with Pto DC3000Q^-^(AvrRpm2_Psa_). Proteins were extracted as in **(B). F**. AtRIN4 proteins (AtRIN4^WT^, AtRIN4^E15AE156A^, AtRIN4^E15A^, and AtRIN4^E156A^) were co-expressed with AvrRpm2_Psa_and RPM1 in *N, benthamiana* to assess HR. The visual inspection (left) and the fluorescence test with the ChemiDoc™ (right) are shown. Red circles denote HR and white circles denote no HR.

As AtRIN4^8A^_E15E156_ recovered ADP-ribosylation from AtRIN4^10A^, the target residue(s) for the AvrRpm2_Psa_ activity could be either E15, E156, or both residues. We created the double mutant allele AtRIN4^E15AE156A^, which is the reciprocal modification of AtRIN4^8A^_E15E156_, as well as the two single mutant alleles AtRIN4^E15A^ and AtRIN4^E156A^. They were transiently expressed in *N. benthamiana* via Agrobacterium and the leaves were infected with Pto DC3000Q^-^(AvrRpm2_Psa_) as above. Western blot analysis showed that AtRIN4^E15AE156A^ was not ADP-ribosylated as expected (Figure 4F). Out of the two individually modified proteins, AtRIN4^E156A^ was not ADP-ribosylated while AtRIN4^E15A^ was still modified, demonstrating that the glutamate (E156) in the C-NOI domain is the target site for AvrRpm2_Psa_ during infection. When the AtRIN4 proteins were co-expressed with AvrRpm2_Psa_ and RPM1 in *N. benthamiana* via Agrobacterium, the RPM1-mediated HR was lost with AtRIN4^E156A^ (Figure 4G, white circle) while the corresponding HR was still detected with AtRIN4^E15A^ (Figure 4G, red circles). Consistent with the earlier observation (Figure 1B), slower migration of ADP-ribosylated AtRIN4 during electrophoresis was detected in AtRIN4^E15A^ or AtRIN4^WT^, but not in AtRIN4^E156A^ or AtRIN4^E15AE156A^ (Figure 4F).

### Conserved glutamate residues in GmRIN4c and GmRIN4d are ADP-ribosylated by AvrRpm2_Psa_

There are four soybean RIN4 homologs (Figure 5A). When co-expressed with AvrRpm1_Pma_ in *N. benthamiana* via Agrobacterium, GmRIN4a and GmRIN4b were shown to be ADP-ribosylated while GmRIN4c or GmRIN4d were not modified (Redditt et al., 2019). In stark contrast, we found that GmRIN4c and GmRIN4d were ADP-ribosylated by Pto DC3000Q^-^ (AvrRpm2_Psa_) while GmRIN4a and GmRIN4b were not modified (Figure 5B). Accordingly, when the GmRIN4 homologs were co-expressed with AvrRpm2_Psa_ and RPM1 in *N. benthamiana* via Agrobacterium, only GmRIN4c and GmRIN4d activated RPM1 (Figure 5C). The two homologs GmRIN4c and GmRIN4d have glutamate (E) residues at their respective positions corresponding to the E156 of AtRIN4. In contrast, GmRIN4a and GmRIN4b have valine (V) at these positions. Similarly, among the two RIN4 homologs of snap bean, PvRIN4b has the glutamate at the position corresponding to the E156 of AtRIN4 while PvRIN4a has valine at that position (Figure 3C). PvRIN4b was ADP-ribosylated with the infection of Pto DC3000Q^-^(AvrRpm2_Psa_) (Figure 3E) and also activated RPM1 when co-expressed with AvrRpm2_Psa_ and RPM1 in *N. benthamiana* (Figure 3F). In contrast, PvRIN4a was not modified by Pto DC3000Q^-^(AvrRpm2_Psa_) and no corresponding HR was detected when co-expressed with AvrRpm2_Psa_ and RPM1 in *N. benthamiana* via Agrobacterium. Therefore, the allelic variations in the critical residue within the C-NOI domains of soybean and snap bean RIN4 homologs were reflected in the differential ADP-ribosylation patterns by AvrRpm2_Psa_, suggesting the conserved glutamate residues at the corresponding positions of the E156 of AtRIN4 are the likely target sites in these RIN4 homologs. Interestingly, the alternative residue valine is also found at the positions corresponding to E156 in other Arabidopsis NOI domain-containing proteins (Redditt et al., 2019).

**Figure 5.**
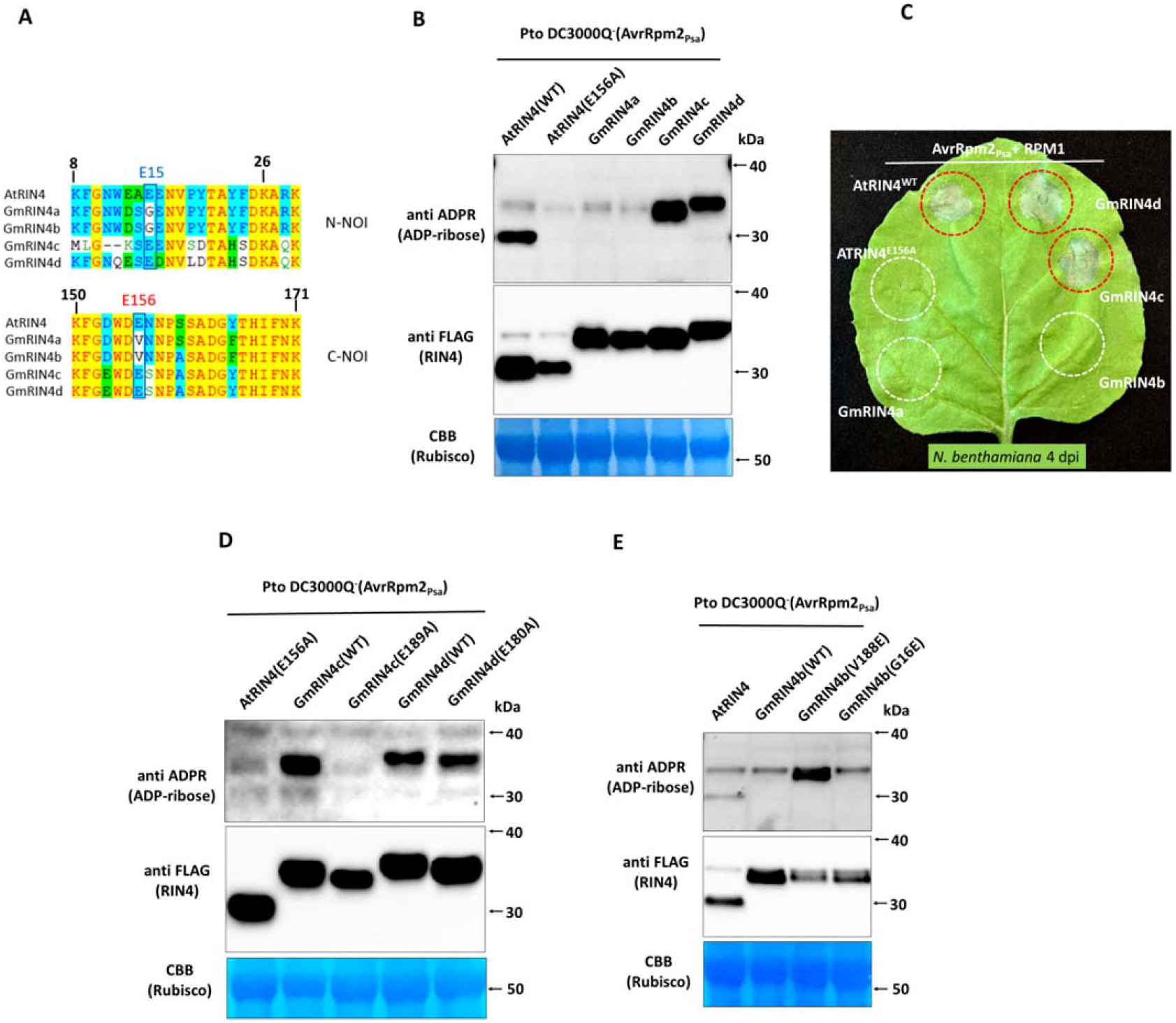
Western blot analysis of soybean RIN4 homologs from *N. benthamiana* leaves infected with Pto DC3000Q^-^(AvrRpm2_Psa_). **A**. Sequence alignment of the nitrate-induced (NOI) sequences in RIN4 homologs of Arabidopsis (AtRIN4) and soybean (GmRIN4a, GmRIN4b, GmRIN4c, and GmRIN4d). E156 in AtRIN4 corresponds to E189 in GmRIN4c or E180 in GmRIN4d, respectively. GmRIN4a and GmRIN4b have a valine (V) at the corresponding positions. **B**. Western blot analysis of soybean RIN4 homologs after infection with Pto DC3000Q^-^(AvrRpm2_Psa_). Proteins were extracted from *N. benthamiana* leaves 1 d post infiltration of Pto DC3000Q^-^(AvrRpm2_Psa_) following Agrobacterium infiltration 2 d earlier for transient expression RIN4 homologs. **C**. Soybean RIN4 homologs were co-expressed with AvrRpm2_Psa_ and RPM1 in *N. benthamiana* via Agrobacterium to assess the RPM1-mediated HR. The visual image was taken in 4 d post Agrobacterium infiltration. Red circles denote HR and white circles denote no HR. **D**. Western blot analysis of GmRIN4c^E189A^ and GmRIN4d^E180A^ from *N. benthamiana* leaves infected with Pto DC3000Q^-^ (AvrRpm2_Psa_). Proteins extraction and Western blot analysis were performed as in (**B**). **E**. Western blot analysis of GmRIN4b^V188E^ and GmRIN4b^G16E^ from *N. benthamiana* leaves infected with Pto DC3000Q^-^(AvrRpm2_Psa_). Proteins extraction and Western blot analysis were performed as in (**B**).

To further study the functional significance of the glutamate residues in the two GmRIN4 homologs at the corresponding positions of E156 in AtRIN4 (E189 in GmRIN4c and E180 GmRIN4d, respectively), substitutions were made to create GmRIN4c^E189A^ and GmRIN4d^E180A^. Proteins were extracted from *N. benthamiana* leaves expressing GmRIN4c^E189A^ or GmRIN4d^E180A^ via Agrobacterium after infection with Pto DC3000Q^-^ (AvrRpm2_Psa_). Western blot analysis showed that GmRIN4c^E189A^ was not ADP-ribosylated, demonstrating the replaced glutamate (E189) was the only target site for AvrRpm2_Psa_ in GmRIN4c (Figure 5D). In contrast, GmRIN4d^E180A^ was still ADP-ribosylated even though the modification of the probable target site was blocked by the substitution (E180A). The ADP-ribosylated GmRIN4d^E180A^ consistently migrated faster than the ADP-ribosylated GmRIN4d^WT^ during electrophoresis (Figure 5D), suggesting potential differences in structure among RIN4 proteins modified at different sites. Unlike GmRIN4c, which has only one target site (E189), GmRIN4d appears to have other residue(s) that can be ADP-ribosylated when E180 is absent.

Apart from the position corresponding to E156 of AtRIN4 in the C-NOI domain, GmRIN4 homologs have another polymorphic site in the corresponding N-NOI domain at the position corresponding to E15 of AtRIN4 (Figure 5A). The soybean RIN4 homologs GmRIN4a and GmRIN4b have substitutions at these two positions when compared with AtRIN4 or the other two GmRIN4 homologs (Figure 5A). To further investigate the participation of these two glutamates in the ADP-ribosylation, we created GmRIN4b^V188E^ and GmRIN4b^G16E^ by instating glutamate residues at the positions corresponding to E156 or E15 of AtRIN4, respectively. Proteins were extracted from *N. benthamiana* leaves transiently expressing the modified GmRIN4b proteins after infection with Pto DC3000Q^-^(AvrRpm2_Psa_). Western blot analysis showed that GmRIN4b^V188E^ became efficiently ADP-ribosylated with a substituted glutamate at the site corresponding to E156 of AtRIN4, while the GmRIN4b^G16E^ with a substituted glutamate corresponding to E15 of AtRIN4 was not modified (Figure 5E). Therefore, the absence of the conserved glutamate residue corresponding to E156 of AtRIN4 is likely responsible for the lack of ADP-ribosylation in GmRIN4b by the bacterial effector AvrRpm2_Psa_.

### A conserved glutamate of MdRIN4-2 is ADP-ribosylated by AvrRpm2_Psa_ and the N-NOI domain plays a role in HR

There are two RIN4 homologs in apple species. In contrast to GmRIN4 homologs, both MdRIN4 homologs have glutamates in the positions corresponding to the E156 of AtRIN4 (E186 in MdRIN4-2 and E184 in MdRIN4-1, Figure 6A). To see whether the E186 in MdRIN4-2 is also the target of AvrRpm2_Psa_, we created MdRIN4-2^E186A^ to block the modification at this site. Proteins were extracted from *N. benthamiana* leaves transiently expressing MdRIN4-2^E186A^ via Agrobacterium after infection with Pto DC3000Q^-^ (AvrRpm2_Psa_). Similar to AtRIN4^E156A^, Western blot analysis showed that MdRIN4-2^E186A^ was not ADP-ribosylated (Figure 6B, left), suggesting that E186 is the likely target of AvrRpm2_Psa_. When MdRIN4-2^E186A^ was co-expressed with AvrRpm2_Psa_ and RPM1 in *N. benthamiana* via Agrobacterium, no significant HR was found compared with the wild type MdRIN4-2^WT^ (Figure 6B, right).

**Figure 6.**
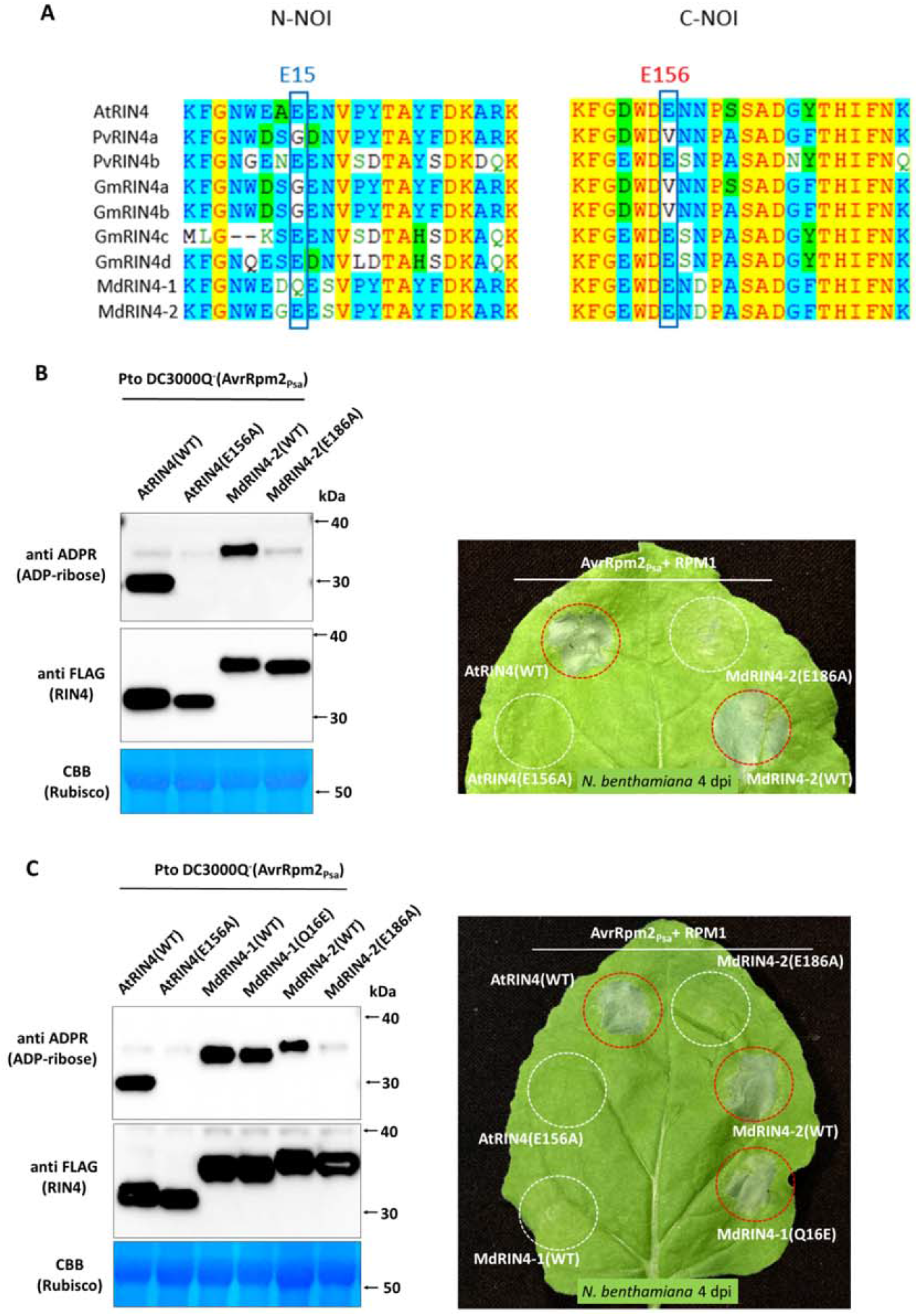
Western blot analysis and RPM1-mediated HR assay of apple RIN4 homologs. **A**. Sequence alignment of the NOI sequences in the RIN4 homologs of Arabidopsis (AtRIN4), snap bean (PvRIN4a and PvRIN4b), soybean (GmRIN4a to GmRIN4d), and apple (MdRIN4-1 and MdRIN4-2). Both apple RIN4 loci have glutamate residues at the positions (E186 in MdRIN4-2 and E184 in MdRIN4-1) corresponding to E156 of AtRIN4. MdRIN4-2 also has a glutamate (E16) at the position corresponding to E15 of AtRIN4, while MdRIN4-1 has an alternative residue glutamine (Q16). **B**. Western blot analysis of MdRIN4-2^E186A^ (left). Proteins were extracted 1 d post infection of Pto DC3000Q^-^(AvrRpm2_Psa_) into *N. benthamiana* leaves that had been pre-infiltrated with Agrobacterium 2 d earlier for transient expression of RIN4 homologs. Apple RIN4 homologs were co-expressed with AvrRpm2_Psa_ and RPM1 in *N. benthamiana* via Agrobacterium to assess the RPM1-mediated HR (right). **C**. Western blot analysis of MdRIN4-1^Q16E^ with other RIN4 homologs (left). Protein extractions and Western blot analysis were performed as in (**B**). MdRIN4-1^Q16E^ and other RIN4 homologs were co-expressed with AvrRpm2_Psa_ and RPM1 via Agrobacterium in different areas of an *N. benthamiana* leaf to assess the RPM1-mediated HR.

Because the C-NOI domains of the two apple RIN4 homologs are identical, two amino acids 15D and 16Q in the N-NOI domain of MdRIN4-1 are the only different residues in the two NOI sequenences when compared with MdRIN4-2. In particular, MdRIN4-2 has a glutamate (E16) in the N-NOI domain at the position corresponding to E15 in AtRIN4, while MdRIN4-1 has glutamine (Q16) at the corresponding position (Figure 6A). Both homologs were ADP-ribosylated by Pto DC3000Q^-^(AvrRpm2_Psa_) as expected (Figure 6C, left). However, when co-expressed with AvrRpm2_Psa_ and AtRIN4 in *N. benthamiana* via Agrobacterium, MdRIN4-1 did not trigger the RPM1-mediated HR, suggesting that the ADP-ribosylation of MdRIN4-1 by AvrRpm2_Psa_ may not be recognised by RPM1 (Figure 6C, right). We created MdRIN4-1^Q16E^ to substitute a glutamate in place of Q16 and tested whether the glutamate at this position would affect the activity. Proteins were extracted from *N. benthamiana* leaves after infection with Pto DC3000Q^-^(AvrRpm2_Psa_) following transient expression of MdRIN4-1^Q16E^ via Agrobacterium. Western blot analysis showed no significant changes in ADP-ribosylation or protein accumulation of MdRIN4-1^Q16E^ compared with the wild type MdRIN4-1^WT^ (Figure 6C, left). However, when MdRIN4-1^Q16E^ was co-expressed with RPM1 and AvrRpm2_Psa_ in *N. benthamiana* via Agrobacterium, an RPM1-mediated HR was observed (Figure 5C, right). The substitution of the glutamate in MdRIN4-1^Q16E^ at the position corresponding to E15 of AtRIN4 resulted in an activation of RPM1, comparable with that of MdRIN4-2. Therefore, even though E16 in MdRIN4-2 (and also in MdRIN4-1^Q16E^) is not modified by AvrRpm2_Psa_, the glutamate residue appears to be also important in activating RPM1. It is interesting to note that apple species also have MdRIN4-1, which is ADP-ribosylated but does not activate RPM1, in addition to the functional MdRIN4-2, which is ADP-ribosylated and activates RPM1. It is possible that the ADP-ribosylation in MdRIN4-1 may be specifically recognised by an apple resistance protein. Alternatively, the host may express a less sensitive RIN4 as well as the fully functional RIN4 to reduce the cost of defence by attenuating unnecessary responses.

### ADP-ribosylation of RIN4 may not be directly linked to phosphorylation

Another effector AvrB from *Pseudomonas syringae* pv. *glycinea* is known to trigger RPM1 activation by inducing phosphorylation at T166 of AtRIN4 (Chung et al., 2011; Lee et al., 2015a). The enzymatic activity of AvrB is not known but the phosphorylation of RIN4 is thought to be mediated by plant kinases such as RIPK (Liu et al., 2011). A previous *in vitro* experiment combined with mass spectrometry analysis suggested that RIPK may have three target residues (T21, S160, and T166) in AtRIN4 (Liu et al., 2011). In particular, the phosphorylation of T166 was proposed as a key physiological switch for defence (Chung et al., 2014). Two other bacterial effectors AvrRpm1_Pma_ and AvrRpm2_Psa_ also trigger the activation of RPM1. In particular, AvrRpm2_Psa_ activates RPM1 by directly ADP-ribosylating E156 of AtRIN4 (Figure 4F and 4G). Therefore, we investigated the impact of the ADP-ribosylation of E156 on the phosphorylation at T166, and *vice versa*, in the RPM1-mediated HR in *N. benthamiana*.

Proteins were extracted from *N. benthamiana* leaves transiently co-expressing RIN4 homologs of Arabidopsis and soybean with AvrB via Agrobacterium. Western blot analysis showed that the RIN4 proteins were not ADP-ribosylated, suggesting that AvrB does not function as an ADP-ribosyl transferase to modify RIN4 (Figure 7A). When the triple mutant AtRIN4^T21AS160AT166A^, in which the potential phosphorylation sites were blocked, was co-expressed with AvrB and RPM1 in *N. benthamiana* via Agrobacterium, there was a significant reduction in the HR compared with AtRIN4^WT^ (Figure 7B, white circle in the right half of the leaf), showing that the AvrB-triggered RPM1 activation depends on these replaced residues as previously reported (Chung et al., 2011). In contrast, when the ADP-ribosylation-deficient allele AtRIN4^E156A^ was co-expressed with AvrB and RPM1, no significant difference was found in the HR compared with the wild type AtRIN4^WT^ (Figure 7B, right half of the leaf). The absence of the ADP-ribosylation site (E156) in AtRIN4^E156A^ had no impact on the AvrB-triggered activation of RPM1. Therefore the AvrB-triggered, phosphorylation-dependent, activation of RPM1 appears not to depend on the ADP-ribosylation of E156 in AtRIN4.

**Figure 7.**
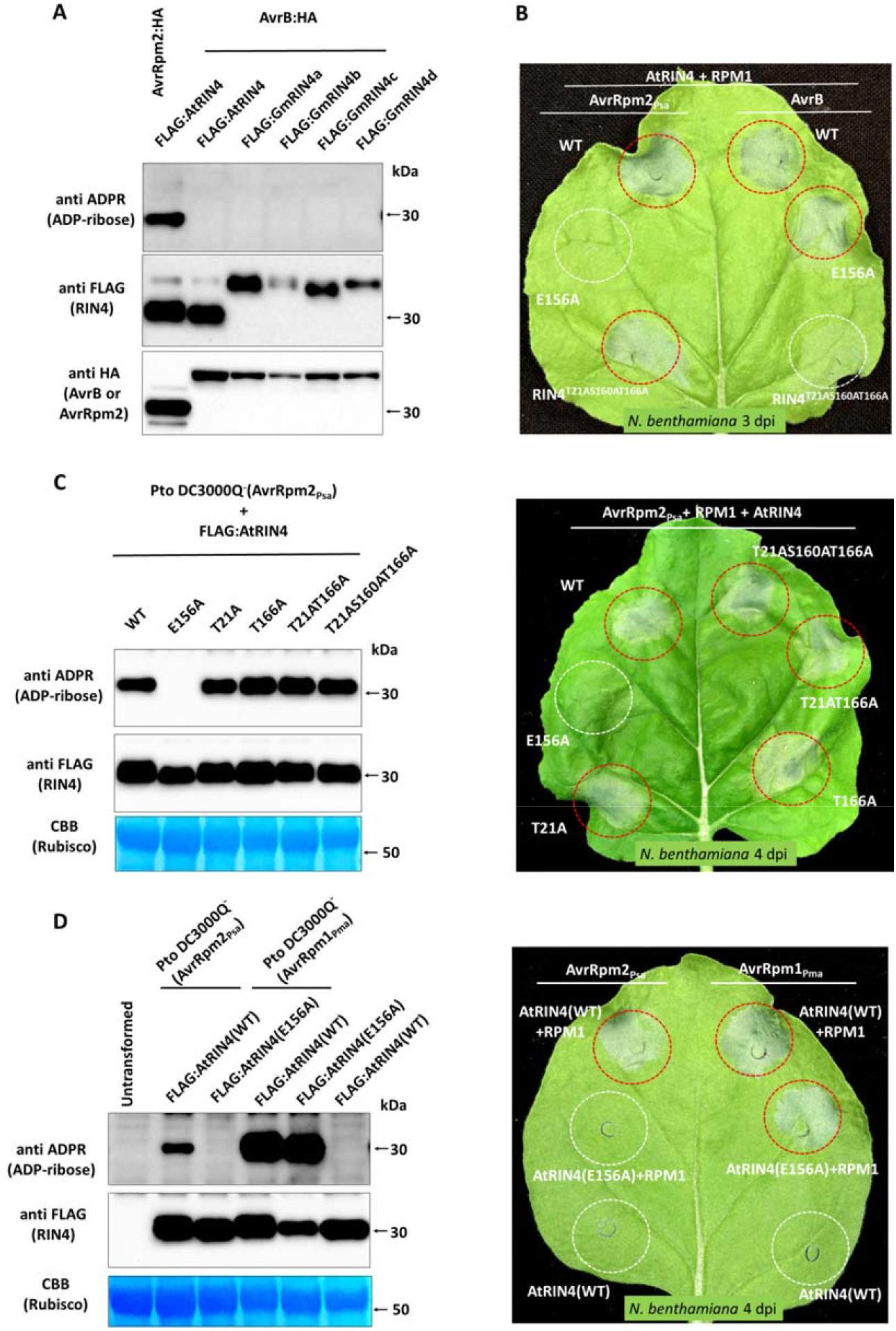
Activities of the type three effectors (T3Es) triggering RPM1 activation. **A**. Western blot analysis of RIN4 homologs of Arabidopsis (AtRIN4) and soybean (GmRIIN4a to GmRIN4d) co-expressed in *N. benthamiana* via Agrobacterium. Proteins were extracted from *N. benthamiana* leaves co-expressing AvrB with RIN4 via Agrobacterium at 2 d post infiltration. **B**. Different AtRIN4 alleles (AtRIN4^WT^, AtRIN4^E156A^ or AtRIN4^T21AS1690AT166A^) and RPM1 were co-expressed with either AvrRpm2_Psa_ (left half) or AvrB (right half) on an *N. benthamiana* leaf via Agrobacterium to assess the corresponding RPM1-mediated HR. **C**. Western blot analysis of different AtRIN4 alleles (AtRIN4^WT^, AtRIN4^E156A^, AtRIN4^T21A^, AtRIN4^T166A^, AtRIN4^T21AT166A^, and AtRIN4^T21AS1690AT166A^) from *N. benthamiana* leaves infected with Pto DC3000Q^-^(AvrRpm2_Psa_) (left). Proteins were extracted 1 d post infection of Pto DC3000Q^-^(AvrRpm2_Psa_) into *N. benthamiana* leaves pre-infiltrated with Agrobacterium 2 d earlier for transient expression of the RIN4 alleles. The AtRIN4 alleles were co-expressed with AvrRpm2_Psa_ and RPM1 on different areas of an *N. benthamiana* leaf via Agrobacterium (right). **D**. Western blot analysis of AtRIN4^E156A^ from *N. benthamiana* leaves infected with either Pto DC3000Q^-^(AvrRpm2_Psa_) or by Pto DC3000Q^-^(AvrRpm1_Pma_) (left). AtRIN4^E156A^ and RPM1 were co-expressed with either AvrRpm2_Psa_ or AvrRpm1_Pma_ on different areas of an *N. benthamiana* leaf to assess the RPM1-mediated HR (right). Red circles denote HR and white circles denote no HR.

In turn, when the phosphorylation-deficient mutant AtRIN4^T21AS160AT166A^ was co-expressed with AvrRpm2_Psa_ and RPM1 in *N. benthamiana* via Agrobacterium, again there was no significant difference in the RPM1-mediated HR compared with the wild type AtRIN4^WT^, while AtRIN4^E156A^ resulted in no host response (Figure 7B, left half of the leaf). Therefore, the AvrRpm2_Psa_-triggered, ADP-ribosylation-dependent, activation of RPM1 appears not to rely on the phosphorylation of those modified sites, including T166. We also tested AtRIN4 alleles with substitutions at the phosphorylation sites in different combinations (AtRIN4^T21A^, AtRIN4^T166A^, AtRIN4^T21AT166A^, and AtRIN4^T21AS160AT166A^) with infection of Pto DC3000Q^-^(AvrRpm2_Psa_) instead of Agrobacterium-mediated transient expression. Western blot analysis showed no significant differences in ADP-ribosylation between AtRIN4^WT^, AtRIN4^T21A^, AtRIN4^T166A^, AtRIN4^T21AT166A^, or AtRIN4^T21AS160AT166A^, while AtRIN4^E156A^ showed no ADP-ribosylation (Figure 7C, left). Accordingly, when the RIN4 alleles were co-expressed with AvrRpm2_Psa_ and RPM1 in *N. benthamiana* via Agrobacterium, all resulted in comparable host responses except AtRIN4^E156A^ (Figure 7C, right).

Even though the two posttranslational modifications, ADP-ribosylation and phosphorylation, of RIN4 are closely linked to the activation of RPM1, they may not be equivalent physiologically as well as biochemically. Because a secreted bacterial effector directly modifies the host protein, ADP-ribosylation of RIN4 by AvrRpm2_Psa_ may be one of the pathogen’s virulence activities. In contrast, the phosphorylation of RIN4 by host kinases is more likely to be a part of defence mechanism responding to a bacterial protein. RPM1 may be independently activated either by the direct ADP-ribosylation of E156 or by the phosphorylation of T166 so long as either modification is recognised by the resistance protein, which is physically associated in the RIN4 multi-protein complex. To our current knowledge, there is no known biochemical activity of AvrB apart from a strong affinity to RIN4. Therefore, AvrB may induce phosphorylation of RIN4 by changing the interactions between proteins, including RIPK and RPM1 in Arabidopsis, in the RIN4 complex. In contrast, AvrRpm2_Psa_ directly modifies RIN4 to generate a distinct structural change, which may affect other physically associated proteins in the multi-protein complex including RPM1 in Arabidopsis.

Apart from the RPM1 activation, we have little knowledge about the physiological significance of the structural change introduced in the RIN4 complex by AvrRpm2_Psa_. It would be interesting to see whether the ADP-ribosylation of RIN4 would affect the activities of other proteins in the RIN4 complex. Interestingly, AvrRpm2_Psa_ also physically associates with RIN4 (Figure 2A), therefore it is possible that the bacterial protein has the ability to change protein-protein interactions in the RIN4 multi-protein complex even without modifying RIN4.

### AvrRpm2_Psa_ and AvrRpm1_Pma_ have distinct properties

The bacterial effector AvrRpm1_Pma_ also functions as an ADP-ribosyl transferase to modify RIN4, leading to the activation of RPM1 (Redditt et al., 2019). To compare the activities of AvrRpm1_Pma_ and AvrRpm2_Psa_, we tested AvrRpm1_Pma_ in combination with AtRIN4^E156A^, which cannot be ADP-ribosylated by AvrRpm2_Psa_. When proteins were extracted from *N. benthamiana* leaves infected with Pto DC3000Q^-^(AvrRpm1_Pma_) following transient expression of AtRIN4^E156A^ via Agrobacterium, Western blot analysis showed that AtRIN4^E156A^ was still ADP-ribosylated, demonstrating that another residue was modified by AvrRpm1_Pma_ (Figure 7D, left). When AvrRpm1_Pma_ was co-expressed with AtRIN4^E156A^ and RPM1 in *N. benthamiana* via Agrobacterium, the corresponding RPM1-mediated HR was detected, while no comparable HR was found with AvrRpm2_Psa_ (Figure 7D, right). AvrRpm1_Pma_ and AvrRpm2_Psa_ also showed completely different activities when tested with GmRIN4 homologs (Figure 5B and Figure S4). Therefore, the activities of these two effectors are not biochemically identical. AvrRpm1_Pma_ and AvrRpm2_Psa_ are the only two ADP-ribosyl transferases (ARTs) from bacterial plant pathogens so far characterised. However, sequence comparisons suggest that ARTs may be widely-distributed among phytopathogens colonising various plant hosts. RPM1 is an Arabidopsis protein and no close homologs are found in most other non-Brassicaceae plant species. Considering most plant species express RIN4 homologs and bacterial ARTs are found in non-Arabidopsis pathogens such as Psa (McCann et al., 2013; Fujikawa and Sawada, 2016; McCann et al., 2017), this posttranslational modification is likely to have other functional significance apart from the RPM1-mediated host response.

### AvrRpm2_Psa_ alleles have differential characteristics in activity and affinity

Based on genetic diversity and toxin production, Psa has been categorized into biovars (bv) (Fujikawa and Sawada, 2016, 2019). Recently bv4 has been transferred to another pathovar, *Pseudomonas syringae* pv. *actinidifoliorum* (Abelleira et al., 2015) and Psa is composed of five bvs currently. Apart from the recently added bv6, all other bvs have AvrRpm2_Psa_ loci. The AvrRpm2_Psa_ alleles in bv1 and bv3 have frameshift mutations, leading to premature termination of the corresponding proteins (Figure 8A). In addition to the frameshift mutations, the AvrRpm2_Psa_ alleles in bv1 and bv3 also have two other independent substitutions when compared with the functional allele in bv5 (AvrRpm2_Psa_^bv5^ or simply AvrRpm2_Psa_). We corrected the frameshifts in the bv1 and bv3 alleles by removing extra nucleotides to express the corresponding full length AvrRpm2_Psa_ proteins. Western blot analysis showed that the edited proteins are expressed in full length (AvrRpm2_Psa_^bv1(in-frame)^ and AvrRpm2_Psa_^bv3(in-frame)^) in *N. benthamiana* (Figure 8B, anti HA in Input panel). However, when the edited proteins were co-expressed with AtRIN4, neither AvrRpm2_Psa_^bv1(in-frame)^ nor AvrRpm2_Psa_^bv3(in-frame)^ modified AtRIN4 (Figure 8B, anti ADPR in Input panel). The frameshift mutations in the bv1 and bv3 alleles are likely to be chronologically the last mutations in these alleles because non-functional genes would only accumulate random mutations. Therefore, the AvrRpm2_Psa_ effectors in bv1 and bv3 appear to have lost their activity to modify RIN4 even before the frameshift mutations were selected.

**Figure 8.**
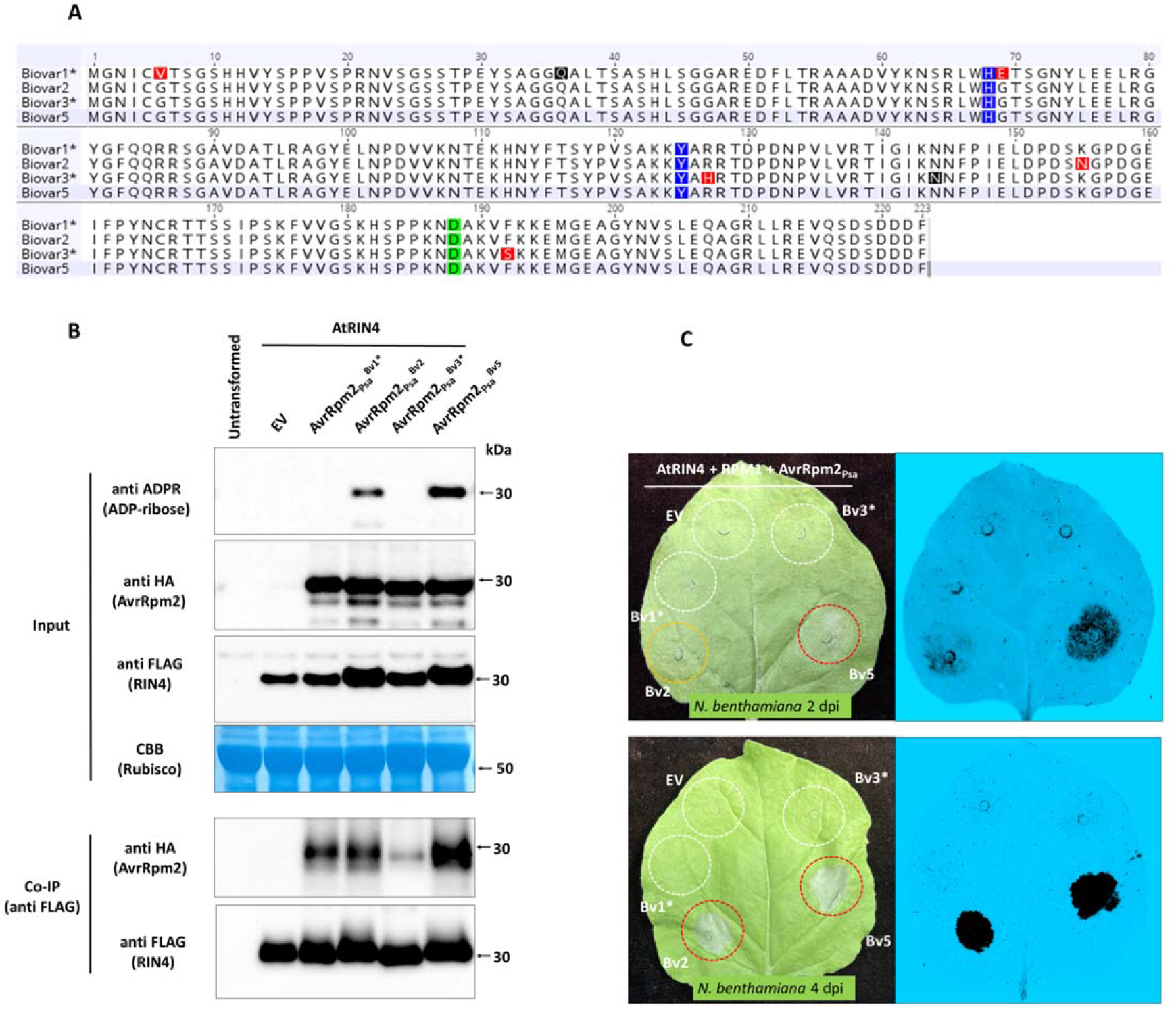
Activity and physical association of AvrRpm2_Psa_ alleles. **A**. Sequence alignment of AvrRpm2_Psa_ alleles from different Psa biovars. The biovar1 and biovar3 alleles were edited to express corresponding full-length proteins by removing extra nucleotides. The corrected sites are marked in black. The two residues H68 and Y125 are marked in blue and D188 is in green. All other substitutions are marked in red. **B**. Western blot analysis of AtRIN4 co-expressed with different AvrRpm2_Psa_ alleles (input panel). Proteins were extracted 2 d after co-infiltration of AtRIN4 and AvrRpm2_Psa_ in *N. benthamiana* via Agrobacterium. Co-IP of AtRIN4 with different AvrRpm2_Psa_ proteins (Co-IP panel). Proteins were immunoprecipitated using anti FLAG magnetic beads to isolate AtRIN4. The precipitated proteins were probed using corresponding antibodies. **C**. AvrRpm2_Psa_ alleles from different Psa biovars were co-expressed with AtRIN4 and RPM1 via Agrobacterium on different areas of an *N. benthamiana* leaf to assess the RPM1-mediated HR. Red circles denote HR, white circles denote no HR, and orange circle denotes reduced HR. Visual inspection (left) and fluorescence detection monitored in the ChemiDoc™ system (right) were shown.

Co-IP analysis with AtRIN4 showed that AvrRpm2_Psa_ proteins from different bvs physically associate with AtRIN4 with differential affinities (Figure 8B, Co-IP panel). In particular, the physical association of AtRIN4 with AvrRpm2_Psa_^bv3(in-frame)^ was significantly lower when compared with AvrRpm2_Psa_^bv5^. The two other substitutions in AvrRpm2_Psa_^bv3(in-frame)^ are R127H and F192S, close to residues Y125 and D188, respectively, which correspond to the two residues in the conserved H-Y-D motif in AvrRpm1_Pma_ (Cherkis et al., 2012). Earlier we found that AvrRpm2_Psa_^Y125A^ lost its catalytic activity while AvrRpm2_Psa_^D188A^ lost the ability for physical association with RIN4 (Figure 2C). Therefore the current AvrRpm2_Psa_^bv3^ allele may have undergone multiple selections to change its ability to affect the RIN4 protein complex (either by ADP-ribosylation of RIN4 or by changing physical association with RIN4). Similarly, the bv1 allele (AvrRpm2_Psa_^bv1^) has two other substitutions in addition to the frameshift mutation when compared with the functional AvrRpm2_Psa_^bv5^. One substitution is G69E, which is next to H68, another residue in the H-Y-D motif. The other substitution in the AvrRpm2_Psa_^bv1^ is G6V. Glycine residues located near N-termini of bacterial effectors are often involved in localisation of the bacterial proteins to membranes and this substitution may have changed the localisation of the bacterial effector in the host.

The main activities of AvrRpm2_Psa_^bv1(in-frame)^ or AvrRpm2_Psa_^bv3(in-frame)^ may not have been the ADP-ribosylation of RIN4. Otherwise, additional frameshift mutations would not have been selected for these loci, which presumably had lost the activities already. AvrRpm2_Psa_^bv1(in-frame)^ may be able to affect the structure of the RIN4 multi-protein complex with a reduced affinity to RIN4 (Figure 8B, CoIP) even though it does not modify RIN4. In contrast, AvrRpm2_Psa_^bv3(in-frame)^ binds to RIN4 significantly weakly, suggesting that this bacterial protein may affect the RIN4 complex neither by direct modification of RIN4 nor by the physical association with RIN4. It is also possible that AvrRpm2_Psa_^bv3(in-frame)^ may have differential affinities for different RIN4 homologs, specifically affecting protein-protein interactions in different RIN4 complexes without modifying RIN4. Alternatively, considering another bacterial ADP-ribosyl transferase AvrRpm1_Pma_ specifically modified at least ten other Arabidopsis proteins containing NOI domains (Redditt et al., 2019), AvrRpm2_Psa_^bv1(in-frame)^ and AvrRpm2_Psa_^bv3(in-frame)^ may have performed other virulence functions, which were phased out by multiple selections, by targeting other proteins.

The allelic variations in the AvrRpm2_Psa_ loci among bv1, bv3, and bv5 demonstrate the importance of the residues around the H-Y-D motif. In contrast, the allele from bv2 (AvrRpm2_Psa_^bv2^) is catalytically functional with a single substitution (K155N) when compared with AvrRpm2_Psa_^bv5^ (Figure 8B and Figure 8C). When co-expressed with RPM1 and AtRIN4 in *N. benthamiana* via Agrobacterium, this allele triggered significantly weaker (or slower) HR compared with AvrRpm2_Psa_^bv5^ (Figure 8C, orange circle in 2 dpi). Co-IP analysis suggests that AvrRpm2_Psa_^bv2^ physically associates with RIN4 with a reduced affinity compared with AvrRpm2_Psa_^bv5^. The replaced residue N155 is distantly located from the H-Y-D motif. It appears that the N155K substitution did not result in the complete loss of catalytic activity (Figure 8B, anti ADPR in Input) nor the ability to physically associate with RIN4 (Figure 8B, anti HA in Co-IP).

Lastly, the allelic variations in the AvrRpm2_Psa_ loci among modern day Psa biovars suggests that there have been substantial interactions between the pathogen and the host relating to these loci, raising a strong possibility that the Psa effector AvrRpm2_Psa_ is recognised in its natural host kiwifruit.

## METHODS

### Plant material, DNA constructs, bacterial strains, and growth conditions

For Agrobacterium-mediated transient expression, open reading frames of AvrRpm2_Psa_ alleles were PCR amplified from respective Psa strains using a primer pair including a reverse primer that contained a HA tag sequence (AR2atgf1 and AR2HAtgar1) listed below. Amplified fragments were cloned in a binary vector (pART27, pHEX, or pGWB) under the 35S promoter to transform the Agrobacterium strain GV3101. The AvrRpm2_Psa_ allele cloned from Psa biovar 5 was used in all experiments unless specified. For Pto infection, a 1.6 Kb genomic AvrRpm2_Psa_ sequence including the *hrpL* box and the linked *ShcF* locus was PCR-amplified from Psa (biovar 5) using a primer pair gAR2psaf1 and gAR2psar1 (listed below). The amplified fragment was cloned in the broad-range vector pBBR1MCS5 to transform Pto DC3000Q^-^, which has was created by deleting the hopQ1 locus (Yoon and Rikkerink, 2020). Constructs containing the AvrRpm1_Pma_ locus were generated from synthesised DNA (a 947 bp EcoRI-EcoRV fragment from *Pseudomonas syringae* pv. *maculicola* plasmid pFKN, based on the Genbank sequence AF359557) with the set of primers (gAR1f1, AR1atgf1, AR1taar1, and AR1HAtgar1) listed below. Other related alleles required for transient expression or infection analyses were either created using the QuickChange Site-directed Mutagenesis Kit (Agilent Technologies, Santa Clara, CA) or synthesised by Twist Bioscience (South San Francisco, CA). *Nicotiana benthamiana* plants were grown at 22°C in long-day conditions (16 h light and 8 h dark) for both Agrobacterium-mediated transient expression and Pto DC3000Q^-^ infection assays. Agrobacterium was freshly grown in LB with appropriate antibiotics at 28°C on a shaker. Cells were concentrated by centrifugation at 4000 g for 10 min and resuspended in infiltration buffer (10 mM MgCl_2_, 100 μM acetosyringone). The cells were diluted to the appropriate concentrations (OD_600_ = 0.01-0.08) and infiltrated into leaf tissues of 4-to 5-week-old plants using a needleless syringe. For the infection experiment, Pto DC3000Q^-^ was re-streaked on a solid King’s B media and re-grown at 28°C for 1 d. The freshly grown Pto DC3000Q^-^ cells were suspended in 10 mM MgCl_2_ for injection. To detect the cell death response in *N. benthamiana*, the Agrobacterium-infiltrated leaves were visually inspected at 2-5 days post infiltration and/or the fluorescence of the infiltrated leaves were monitored at 1-2 d post infiltration under the Pro-Q Emerald 488 in the ChemiDoc™ Gel Imaging System (Bio-Rad, Hercules, CA, USA) as described previously (Yoon and Rikkerink 2020).

### Agrobacterium-mediated transient expression and pathogen infection in *N. benthamiana*

For all plant transformations, the Agrobacterium strain GV3101 was transformed by electroporation with a binary vector (pART27, pHEX2, or pGWB) containing a specified construct under the 35S promoter. For transient expression, 4-5 weeks old *N. benthamiana* leaves were syringe-infiltrated with Agrobacterium suspension (OD_600_=0.01 to 0.08) carrying the expression construct. A new sequential assay combined the Agrobacterium-mediated transient expression of a host protein with pathogen infection to deliver bacterial effectors with the T3SS. In such an assay *N. benthamiana* leaves were first infiltrated with Agrobacterium, carrying the specified plant geneexpression construct, which was freshly grown in LB media and adjusted to OD_600_=0.04. Two d post Agrobacterium infiltration, the pre-infiltrated *N. benthamiana* leaves were infected using a needleless syringe with a Pto strain DC3000 without hopQ1 (Pto DC3000Q^-^) (Wei et al., 2007) (Yoon and Rikkerink, 2020), carrying the bacterial effector. The Pto suspension was derived from freshly grown Pto in King’s B media and adjusted to OD_600_=1.0. Leaf discs were collected at 15-24 h post Pto infection for protein extraction before leaves collapsed due to the progression of the infection.

### Western blot analysis and co-immunoprecipitation

Western blot analysis and co-immunoprecipitation (Co-IP) were performed as previously described with some modifications (Yoon and Rikkerink, 2020). For immunoblot analysis, two leaf discs (8 mm diameter) were collected from infiltrated areas of the leaves and placed in microfuge tubes containing cold 250 µL of extraction buffer (2% SDS, 10% glycerol in PBS). Tissues were homogenised on ice using a micro-pestle and incubated at 90°C for 8 min and centrifuged at 16000 g for 2 min to collect the supernatant. Aliquots were run on a 4–12% SDS-PAGE gradient gel and electrophoresed proteins were transferred to a PDVF membrane (Immobilon-P, Millipore, Burlington, MA, USA). To facilitate detection, an in frame C-terminal HA tag was added onto effector sequences (avrRpm2_Psa_, avrRpm1_Pma_ or AvrB) and an in frame N-terminal FLAG tag was added onto RIN4 homolog plant sequences. Primary antibodies were used at a 1:5000 dilution and an anti-mouse secondary antibody (A9044, Sigma-Adrich) was used at a 1:10 000 dilution. For Co-IP analyses, Agrobacterium-infiltrated *N. benthamiana* tissues were collected 2 d post infiltration. A 0.5 g sample of the collected tissues were ground under liquid nitrogen and suspended in 1 ml of co-IP extraction buffer (1× PBS, 1% n-dodecyl-β-d-maltoside (Invitrogen, Carlsbad, CA), protease inhibitor cocktail cOmplete™ (Sigma-Aldrich, St Louis, MO) in NativePAGE™ buffer (Invitrogen)). Extracted protein samples were centrifuged at 10000 g for 2 min and the supernatant was collected. After IP using the Dynabeads™ Protein G Immunoprecipitation Kit (1007D, Invitrogen), Western blots were prepared and probed using HRP-conjugated primary antibodies in 0.2% I-Block (T2015, Invitrogen). The antibodies used were F1804 (anti-FLAG, Sigma-Aldrich St Louis, MO, USA), A8592 (anti-FLAG-HRP, Sigma-Aldrich), H9658 (anti-HA, Sigma-Aldrich), 12013819001 (3F10, anti-HA-HRP, Roche, Basel, Switzerland), and SAB4700447 (anti-Myc, Sigma-Aldrich) at 1:5000 (v/v) dilutions. To detect ADP-ribosylated proteins, 1:5000 (v/v) dilution of an anti-ADPR reagent (MABE1016, EMD Millipore) was used with the anti-rabbit secondary antibody (A0545, Sigma-Aldrich). Proteins were visualized using the ECL Clarity or the ECL Clarity Max detection systems (Bio-Rad). The developed Western blots were rinsed with PBS and post-stained with Coomassie Brilliant Blue (CBB) to visualise protein bands for normalising protein loading.

### Mass Spectrometry analysis

Four to five weeks old *N. benthamiana* leaves were syringe-infiltrated with a mixture of freshly grown Agrobacterium cultures of FLAG:AtRIN4 and AvrRpm1_Pma_ (or AvrRpm2_Psa_) in equal amounts (OD_600_= 0.2 each) in the infiltration buffer (10 mM MgCl_2_, 100 μM acetosyringone). At 2 d post infiltration, 1 g of infiltrated leaves were ground under liquid nitrogen to a fine powder. Two mL of extraction buffer (Pearce Co-IP buffer, Thermo-Fisher) was added to the ground tissues and the resulting suspension was thoroughly mixed. The suspended mix was left at room temperature for 30 min with agitation and then centrifuged at 10000 rpm for 2 min to collect the supernatant. A suspension of 100 μL of anti FLAG-conjugated magnetic beads (Pearce, Thermo-Fisher) was added to the collected fraction and the mixture was left at room temperature for 30 min with agitation. After this precipitation, the magnetic beads with immuno-precipitated proteins were collected and washed four times with PBS. 50 μL of SDS-PAGE sample buffer was added to the washed magnetic beads with attached proteins and the proteins were extracted from the magnetic beads by heating the mix at 55°C for 12 min. The extracted proteins were run on a 4-12% SDS-PAGE gradient gel in MOPS running buffer. When the electrophoresis was completed, the gel was stained with Coomassie Brilliant Blue (CBB) and the visualised RIN4 protein bans was excised from the gel for mass spectrometry analysis.

Gel bands were destained, dehydrated with acetonitrile, and then subjected to reduction with 5 mM dithiothreitol, alkylation with 15 mM iodoacetamide, and digestion with 12.5 ng/μl modified sequencing grade porcine trypsin (Promega, Madison, WI) at 45°C for 1 hour. The digest was acidified with formic acid and then injected onto a 0.3×10 mm trap column packed with Reprosil C18 media (ESI Source Solutions, Woburn, MA) and desalted for 5 minutes at 10 μl/min before being separated on a 0.075 × 200 mm picofrit column (New Objective, Littleton, MA) packed with 3 μm Reprosil C18 media. The following gradient was applied at 300 nl/min using an Eksigent NanoLC 425 UPLC system (AB SCIEX, Concord, ON): 0 min 5% B; 20 min 40% B; 22 min 98% B; 24 min 98% B; 25 min 5% B; 30 min 5% B, where B was 0.1% formic acid in acetonitrile and the remainder of these solutions was solution A (0.1% formic acid in water). The picofrit spray was directed into a TripleTOF 6600 Quadrupole-Time-of-Flight mass spectrometer (AB SCIEX) scanning from 300-2000 m/z for 200ms, followed by 40ms MS/MS scans on the 35 most abundant multiply-charged peptides (m/z 80-1600). The mass spectrometer and HPLC system were under the control of the Analyst TF 1.8 software package (AB SCIEX). The resulting MS/MS data was searched against an in-house database comprising a set of common contaminant sequences, the FLAG-tagged Arabidopsis RIN4 sequence, plus *Nicotiana* and Rat entries from Uniprot.org (183,885 entries in total), using ProteinPilot version 5.0 (AB SCIEX). The peptide summary exported from ProteinPilot was further processed in Excel using a custom macro to remove proteins with Unused Scores below 1.3, eliminate inferior or redundant peptide spectral matches, and to sum the intensities for all unique peptides from each protein.

### Accession numbers

Sequence data from this article can be found in the EMBL/GenBank data libraries under the following accession number(s): AtRIN4 (AT3G25070; NP_001325873), GmRIN4a (Glyma03g19920; NP_001235221), GmRIN4b (Glyma16g12160; NP_001239973), GmRIN4c (Glyma18g36000; NP_001235235), GmRIN4d (Glyma08g46400; NP_001235252), PvRIN4a (XP_007134125.1), PvRIN4b (XP_007140654.1), MdRIN4-1(NM_001293994.1), MdRIN4-2 (NP_001280834.1), AvrB (WP_122390765.1), AvrRpm1_Pma_ (NP_114197), AvrRpm2_Psa_ (WP_099978761.1), RPM1 (At3g07040).

### Primers used in this study

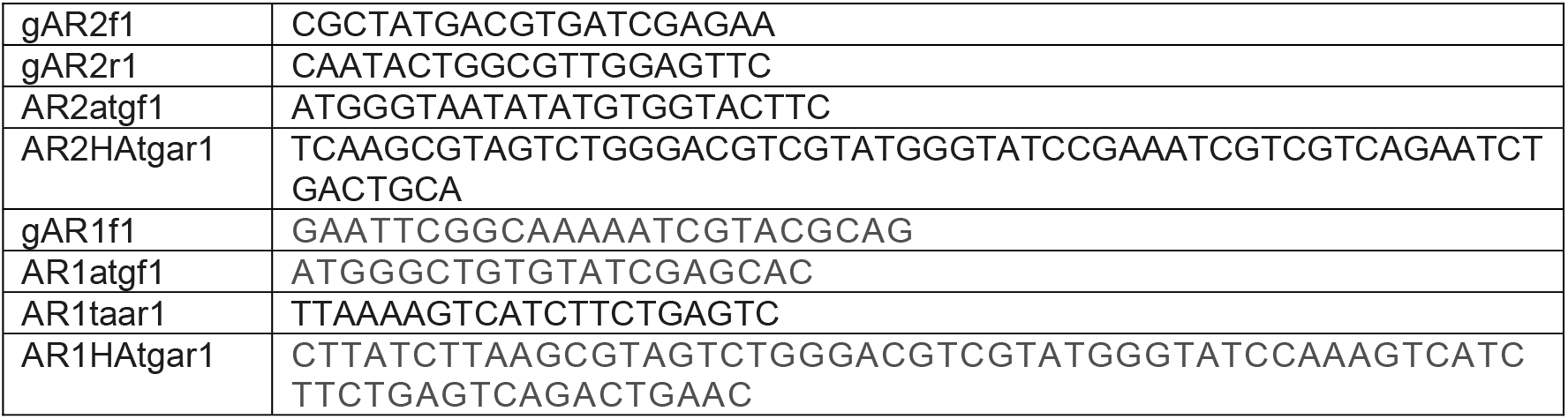

## SUPPLEMENTAL DATA

**Fig S1**. LC-MS/MS analysis of AtRIN4 co-expressed with AvrRpm1_Pma_ in *N. benthamiana* via Agrobacterium.

**Fig S2**. Western blot analysis of modified RIN4 transiently co-expressed with AvrRpm1_Pma_ in *N. benthamiana* via Agrobacterium.

**Fig S3**. LC-MS/MS analysis of AtRIN4 co-expressed with AvrRpm2_Psa_ in *N. benthamiana* via Agrobacterium.

**Fig S4**. Western blot analysis of soybean RIN4 homologs modified by Pto DC3000Q^-^ (AvrRpm1_Pma_).

## ACKNOWLEDGEMENT

The authors thank Prof Andrew Allan and Dr Joanna Bowen of the New Zealand Institute for Plant and Food Research for critically reviewing the manuscript. This work was supported by the Plant and Food Research Fund KRIP PSA & RIN4 (P/880002/13). Further experimental details will be available upon request.

## AUTHOR CONTRIBUTIONS

MY and ER conceived the study. MY designed and conducted experiments, and wrote the manuscript with input from all authors. MM conducted mass spectrometry analysis and wrote the mass spectrometry section in the Methods. ER acquired research funding, contributed to research through discussions, and edited the manuscript.

## FIGURE LEGENDS

**Figure 1**. Transient co-expression of AtRIN4 and AvrRpm2_Psa_ (biovar 5) in *N. benthamiana* via Agrobacterium. **A**. Sequence alignment of AvrRpm2_Psa_ and AvrRpm1_Pma_. The residues (H63, Y122, and D185) in the conserved H-Y-D motif of AvrRpm1_Pma,_ proposed previously (Cherkis et al., 2012), are marked with blue labels. The corresponding residues in the H-Y-D motif of AvrRpm2_Psa_ are H68, Y125, and D188. **B**. Western blot analysis demonstrating the ADP-ribosyl transferase activity of AvrRpm2_Psa_. Proteins were extracted from *N. benthamiana* leaves co-expressing AvrRpm2_Psa_:HA and FLAG:AtRIN4 via Agrobacterium at 2 d post infiltration. ADP-ribosylated proteins were detected using the anti ADPR binding reagent. FLAG:RIN4 and AvrRpm2_Psa_:HA were detected using corresponding antibodies. **C**. RPM1-mediated HR (hypersensitive response) assay. An *N. benthamiana* leaf co-expressing FLAG:AtRIN4 and RPM1:Myc with either AvrRpm1_Pma_ or AvrRpm1_Pma_ in different combinations (1: AvrRpm1_Pma_^WT^:HA (wild type); 2: AvrRpm1_Pma_^3A^ (triple substitutions: H63A, Y122A, D185A); 3: AvrRpm2_Psa_^WT^:HA (wild type); 4: AvrRpm2_Psa_^3A^ (triple substitutions: H68A, Y125A, D188A) delivered via Agrobacterium. Red circles denote HR and white circles denote no HR. The fluorescence of the same leaf monitored under a 488 nm tray in the ChemiDoc™ is also shown (bottom). Images were taken 2 d post Agrobacterium infiltration. **D**. Western blot analysis using proteins extracted from the corresponding areas of the leaf used in **C** 2 d post Agrobacterium injection.

**Figure 2**. Western blot analysis of AtRIN4 transiently co-expressed with AvrRpm2_Psa_ in *N. benthamiana* via Agrobacterium. **A**. Proteins were extracted from *N. benthamiana* leaves co-expressing FLAG:AtRIN4 with either Empty vector (lane 1), HA:AtRIN4 (lane 2), or AvrRpm2_Psa_:HA (lane 3) via Agrobacterium. Protein extracts were immunoprecipitated using anti HA magnetic beads. Precipitated proteins were probed with corresponding antibodies. **B**. Western blot analysis showed the ADP-ribosylated AtRIN4 migrated more slowly compared with unmodified AtRIN4 during SDS-PAGE. Protein extracts from *N. benthamiana* leaves co-expressing FLAG:AtRIN4 with AvrRpm2_Psa_:HA (even-numbered) or expressing only FLAG:AtRIN4 (odd-numbered) were probed. **C**. Western blot analysis of different AvrRpm2_Psa_ alelles (AvrRpm2_Psa_^WT^, AvrRpm2_Psa_^H68A^, AvrRpm2_Psa_^Y125A^, AvrRpm2_Psa_^H188A^, and AvrRpm2_Psa_^3A^) co-expressed with AtRIN4 via Agrobacterium (Input panel). For Co-IP of modified AvrRpm2 with AtRIN4 (Co-IP panels), protein extracts were precipitated using anti FLAG magnetic beads to isolate FLAG:AtRIN4. Precipitated proteins were probed using corresponding antibodies. **D**. Modified AvrRpm2 alleles (AvrRpm2_Psa_^H68A^, AvrRpm2 _Psa_^Y125A^, and AvrRpm2_Psa_^D188A^) were co-expressed with RPM1 and AtRIN4 on an *N. benthamiana* leaf to assess the corresponding RPM1-mediated HR. **E**. Combinations of modified AvrRpm2 alleles (AvrRpm2_Psa_^D188A^ with AvrRpm2_Psa_^H68A^ and AvrRpm2_Psa_^D188A^ with AvrRpm2_Psa_^Y125A^) were co-expressed with RPM1 and AtRIN4 on an *N. benthamiana* leaf. Red circles denote HR and white circles denote no HR. Images were taken at 3 d post Agrobacterium injection (**D** and **E**)

**Figure 3**. Secretion of bacterial effector via infection of Pto DC3000Q^-^ in *N. benthamiana*. **A**. An *N. benthamiana* leaf was infected with Pto DC3000Q^-^(AvrRpm2_Psa_) at a high concentration (OD_600_=1.0, or 5×10^8^ cfu/mL) in the area pre-infiltrated with Agrobacterium 2 d earlier for transient expression of AtRIN4 (marked with a circle, bottom). Dark discolouration and leaf margin malformations indicate Pto infection. The control area infiltrated only with Agrobacterium is also marked (top). Image was taken at 15 h post infection. **B**. Western blot analysis demonstrated that AtRIN4 was ADP-ribosylated upon infection with Pto DC3000Q^-^(AvrRpm2_Psa_). Proteins were extracted from *N. benthamiana* leaves expressing AtRIN4 via Agrobacterium after infection with Pto DC3000Q^-^ (AvrRpm2_Psa_). ADP-ribosylated proteins were detected using anti ADPR binding reagent. (EV: control Pto DC3000Q^-^ with an empty vector). **C**. Alignment of the two NOI sequences in RIN4 homologs of Arabidopsis (AtRIN4), snap bean (PvRIN4a and PvRIN4b) and soybean (GmRIN4a to GmRIN4d). **D**. Western blot analysis of RIN4 co-expressed with AvrRpm2_Psa_ *in planta* via Agrobacterium. **E**. Western blot analysis of RIN4 homologs infected with Pto DC3000Q^-^(AvrRpm2_Psa_). Proteins were extracted as in (**B**). **F**. The indicated RIN4 homologs were co-expressed with AvrRpm2_Psa_ and RPM1 in different areas of an *N. benthamiana* leaf via Agrobacterium to assess the corresponding RPM1-mediated HR. Red circles denote HR and white circles denote no HR.

**Figure 4**. Identification of the target residue for AvrRpm2_Psa_ by mutational analysis. **A**. Modifications generated in the two NOI sequences of AtRIN4. The ten modifications (N11A, E13A, E15A, E16A, N17A, D153A, D155A, E156A, N157A, N158A) in AtRIN4^10A^ are boxed in black () and the six modifications in AtRIN4^6A^ (N11A, E16A, N17D, D153A, N157A, N158A) are marked with asterisk (* in N11 and D153, previously identified by Redditt et al.) (Redditt et al., 2019) or closed circle (• in E16, N17, N157, and N158 identified in this study). **B**. Western blot analysis of RIN4 proteins (AtRIN4^WT^, GmRIN4b^WT^, AtRIN4^6A^ and AtRIN4^10A^) from leaves infected with with Pto DC3000Q^-^(AvrRpm2_Psa_). Proteins were extracted 1 d post infection of Pto DC3000Q^-^(AvrRpm2) in *N. benthamiana* leaves pre-infiltrated with Agrobacterium 2 d earlier for transient expression of RIN4. **C**. AtRIN4^10A^ was co-expressed with AvrRpm2_Psa_ and RPM1 in *N. benthamiana* to assess the RPM1-mediated HR. HR was assessed visually (left) or by monitoring fluorescence with ChemiDoc™ (right) at 2 d post Agrobacterium infiltration. **D**. Western blot analysis of the five AtRIN4^8A^ variant proteins (AtRIN4^8A^_E11D153_, AtRIN4^8A^_E13D155_, AtRIN4^8A^_E15E156_, AtRIN4^8A^_E16N157_, and AtRIN4^8A^_N17N158_) from leaves infected with with Pto DC3000Q^-^ (AvrRpm2_Psa_). Proteins were extracted as in (**B). E**. The five AtRIN4^8A^ proteins were co-expressed with AvrRpm2_Psa_ and RPM1 in *N. benthamiana* via Agrobacterium to assess the corresponding HR. Visual inspection (left) and fluorescence test using the ChemiDoc™ (right) are shown. **F**. Western blot analysis of AtRIN4 proteins (AtRIN4^WT^, AtRIN4^E15AE156A^, AtRIN4^E15A^, and AtRIN4^E156A^) from leaves infected with Pto DC3000Q^-^(AvrRpm2_Psa_). Proteins were extracted as in (**B). G**. AtRIN4 proteins (AtRIN4^WT^, AtRIN4^E15AE156A^, AtRIN4^E15A^, and AtRIN4^E156A^) were co-expressed with AvrRpm2_Psa_ and RPM1 in *N. benthamiana* to assess the RPM1-mediated HR. The visual inspection (left) and the fluorescence test with the ChemiDoc™ (right) are shown. Red circles denote HR and white circles denote no HR.

**Figure 5**. Western blot analysis of soybean RIN4 homologs from *N. benthamiana* leaves infected with Pto DC3000Q^-^(AvrRpm2_Psa_). **A**. Sequence alignment of the nitrate-induced (NOI) sequences in RIN4 homologs of Arabidopsis (AtRIN4) and soybean (GmRIN4a, GmRIN4b, GmRIN4c, and GmRIN4d). E156 in AtRIN4 corresponds to E189 in GmRIN4c or E180 in GmRIN4d, respectively. GmRIN4a and GmRIN4b have a valine (V) at the corresponding positions. **B**. Western blot analysis of soybean RIN4 homologs after infection with Pto DC3000Q^-^(AvrRpm2_Psa_). Proteins were extracted from *N. benthamiana* leaves 1 d post infiltration of Pto DC3000Q^-^(AvrRpm2_Psa_) following Agrobacterium infiltration 2 d earlier for transient expression RIN4 homologs. **C**. Soybean RIN4 homologs were co-expressed with AvrRpm2_Psa_ and RPM1 in *N. benthamiana* via Agrobacterium to assess the RPM1-mediated HR. The visual image was taken in 4 d post Agrobacterium infiltration. Red circles denote HR and white circles denote no HR. **D**. Western blot analysis of GmRIN4c^E189A^ and GmRIN4d^E180A^ from *N. benthamiana* leaves infected with Pto DC3000Q^-^ (AvrRpm2_Psa_). Proteins extraction and Western blot analysis were performed as in (**B**). **E**. Western blot analysis of GmRIN4b^V188E^ and GmRIN4b^G16E^ from *N. benthamiana* leaves infected with Pto DC3000Q^-^(AvrRpm2_Psa_). Proteins extraction and Western blot analysis were performed as in (**B**).

**Figure 6**. Western blot analysis and RPM1-mediated HR assay of apple RIN4 homologs. **A**. Sequence alignment of the NOI sequences in the RIN4 homologs of Arabidopsis (AtRIN4), snap bean (PvRIN4a and PvRIN4b), soybean (GmRIN4a to GmRIN4d), and apple (MdRIN4-1 and MdRIN4-2). Both apple RIN4 loci have glutamate residues at the positions (E186 in MdRIN4-2 and E184 in MdRIN4-1) corresponding to E156 of AtRIN4. MdRIN4-2 also has a glutamate (E16) at the position corresponding to E15 of AtRIN4, while MdRIN4-1 has an alternative residue glutamine (Q16). **B**. Western blot analysis of MdRIN4-2^E186A^ (left). Proteins were extracted 1 d post infection of Pto DC3000Q^-^(AvrRpm2_Psa_) into *N. benthamiana* leaves pre-infiltrated with Agrobacterium 2 d earlier for transient expression of RIN4 homologs. Apple RIN4 homologs were co-expressed with AvrRpm2_Psa_ and RPM1 in *N. benthamiana* via Agrobacterium to assess the RPM1-mediated HR (right). **C**. Western blot analysis of MdRIN4-1^Q16E^ with other RIN4 homologs (left). Protein extractions and Western blot analysis were performed as in (**B**). MdRIN4-1^Q16E^ and other RIN4 homologs were co-expressed with AvrRpm2_Psa_ and RPM1 via Agrobacterium in different areas of an *N. benthamiana* leaf to assess the RPM1-mediated HR.

**Figure 7**. Activities of the type three effectors (T3Es) triggering RPM1 activation. **A**. Western blot analysis of RIN4 homologs of Arabidopsis (AtRIN4) and soybean (GmRIIN4a to GmRIN4d) co-expressed in *N. benthamiana* via Agrobacterium. Proteins were extracted from *N. benthamiana* leaves co-expressing AvrB with RIN4 via Agrobacterium at 2 d post infiltration. **B**. Different AtRIN4 alleles (AtRIN4^WT^, AtRIN4^E156A^ or AtRIN4^T21AS1690AT166A^) and RPM1 were co-expressed with either AvrRpm2_Psa_ (left half) or AvrB (right half) on an *N. benthamiana* leaf via Agrobacterium to assess the corresponding RPM1-mediated HR. **C**. Western blot analysis of different AtRIN4 alleles (AtRIN4^WT^, AtRIN4^E156A^, AtRIN4^T21A^, AtRIN4^T166A^, AtRIN4^T21AT166A^, and AtRIN4^T21AS1690AT166A^) from *N. benthamiana* leaves infected with Pto DC3000Q^-^(AvrRpm2_Psa_) (left). Proteins were extracted 1 d post infection of Pto DC3000Q^-^(AvrRpm2_Psa_) into *N. benthamiana* leaves pre-infiltrated with Agrobacterium 2 d earlier for transient expression of the RIN4 alleles. The AtRIN4 alleles were co-expressed with AvrRpm2_Psa_ and RPM1 on different areas of an *N. benthamiana* leaf via Agrobacterium (right). **D**. Western blot analysis of AtRIN4^E156A^ from *N. benthamiana* leaves infected with either Pto DC3000Q^-^(AvrRpm2_Psa_) or by Pto DC3000Q^-^(AvrRpm1_Pma_) (left). AtRIN4^E156A^ and RPM1 were co-expressed with either AvrRpm2_Psa_ or AvrRpm1_Pma_ on different areas of an *N. benthamiana* leaf to assess the RPM1-mediated HR (right). Red circles denote HR and white circles denote no HR.

**Figure 8**. Activity and physical association of AvrRpm2_Psa_ alleles. **A**. Sequence alignment of AvrRpm2_Psa_ alleles from different Psa biovars. The biovar1 and biovar3 alleles were edited to express corresponding full-length proteins by removing extra nucleotides. The corrected sites are marked in black. The two residues H68 and Y125 are marked in blue and D188 is in green. All other substitutions are marked in red. **B**. Western blot analysis of AtRIN4 co-expressed with different AvrRpm2_Psa_ alleles (input panel). Proteins were extracted 2 d after co-infiltration of AtRIN4 and AvrRpm2_Psa_ in *N. benthamiana* via Agrobacterium. Co-IP of AtRIN4 with different AvrRpm2_Psa_ proteins (Co-IP panel). Proteins were immunoprecipitated using anti FLAG magnetic beads to isolate AtRIN4. The precipitated proteins were probed using corresponding antibodies. **C**. AvrRpm2_Psa_ alleles from different Psa biovars were co-expressed with AtRIN4 and RPM1 via Agrobacterium on different areas of an *N. benthamiana* leaf to assess the RPM1-mediated HR. Red circles denote HR, white circles denote no HR, and orange circle denotes reduced HR. Visual inspection (left) and fluorescence detection monitored in the ChemiDoc™ system (right) were shown.

## Supplemental Data

**Figure S1**. LC-MS/MS analysis of AtRIN4 co-expressed with AvrRpm1_Pma_ in *N. benthamiana* via agrobacterium. **A**. A Coomassie Brilliant Blue (CBB) stained SDS-PAGE gel after immunoprecitation of protein extracts from *N. benthamiana* leaves co-infiltrated with FLAGA:AtRIN4 and AvrRpm1_Pma_ via agrobacterium (EV: empty vector). **B**. Confirmation of excised FLAG:AtRIN4 by Western blot analysis using anti FLAG antibody. FLAG:AtRIN4 band was excised from the gel and boiled in 2× sampling buffer for 5 min (IP). **C**. Summary of fragment ions in the LC-MS/MS spectra of the modified AtRIN4 peptides (not all fragmentations are shown). ADP-ribose moieties were identified at N157, N158, and N17. Fragmentation evidence also suggested that either E15 or E16 was modified in one peptide (top). **D**. An example of LC-MS/MS fragmentation evidence, showing N17 was modified with a ribose derived from ADP-ribose. (ADP-R: ADP-ribose, P-R: phospho-ribose, R: ribose, y: y-ions, b: b-ions).

**Figure S2**. Western blot analysis of modified RIN4 transiently co-expressed with AvrRpm1_Pma_ in *N. benthamiana* via Agrobacterium. **A**. Western blot of proteins extracted form *N. benthamiana* leaves co-expressing AvrRpm1_Pma_ with GmRIN4b^N12AD185A^ or AtRIN4^N11AD153A^, modified based on previous mass spectrometry analyses (Redditt et al., 2019), via agrobacterium. **B**. Western blot of proteins extracted form *N. benthamiana* leaves co-expressing AvrRpm1_Pma_ with AtRIN4^E16AN17AN157AN158A^, modified based on our mass spectrometry analysis, or AtRIN4^E11AE16AN17AD153N157AN158A^, with all collectively identified residues blocked, via agrobacterium. **C**. Modified RIN4 homologs (GmRIN4b^N12AD185A^, AtRIN4^N11AD153A^, and AtRIN4^E16AN17AN157AN158A^) and unmodified RIN4 homologs (AtRIN4^WT^ and GmRIN4b^WT^) were co-expressed with RPM1 and AvrRpm1_Pma_ in *N. benthamiana* via agrobacterium. YFP was also co-expressed with RPM1 and AvrRpm1_Pma_ as a negative control for RPM1-mediated response. Red circles denote HR and white circle denotes no HR.

**Figure S3**. LC-MS/MS analysis of AtRIN4 co-expressed with AvrRpm2_Psa_ in *N. benthamiana* via agrobacterium. **A**. Summary of fragment ions in the LC-MS/MS spectra of the modified AtRIN4 peptides. The locations of the targets predicted from the LC-MS/MS fragmentation are marked with blue boxes. Identified ADP-ribose moieties are labelled in orange. **B**. An example of fragmentation evidence suggesting the target site is either D155, E156, N157, or N158 in this peptide. (ADP-R: ADP-ribose, R: ribose, y: y-ions, b: b-ions).

**Figure S4**. Western blot analysis of soybean RIN4 homologs modified by Pto DC3000Q^-^ (AvrRpm1_Pma_). Proteins were extracted from *N. benthamiana* leaves infected with Pto DC3000Q^-^(AvrRpm1_Pma_) in 1 d post infection, following Agrobacterium infiltration 2 d earlier for transient expression of RIN4 homologs.

